# A domain-general neural signature of serial order memory across action and perception

**DOI:** 10.64898/2026.03.27.714833

**Authors:** Alexandros Karagiorgis, Susanne Dyck, Anwesha Das, Katja Kornysheva, Elena Azañón, Max-Philipp Stenner

**Author notes:** Department of Economics, University of Applied Sciences Magdeburg-Stendal, Magdeburg, Germany (Anwesha Das). These authors contributed equally to this work (Elena Azañón, Max-Philipp Stenner).

## Abstract

Remembering events in the correct order, and generating ordered sequences of actions, are fundamental abilities across species. Behavioral studies, and theoretical work, raise the possibility that the brain represents serial order by a domain-general neural code, following the principle of Competitive Queuing. However, direct neurophysiological evidence for Competitive Queuing exists only in the motor domain. When humans and non-human primates prepare for a series of movements, several of the upcoming movements are represented in parallel, with their representational strength reflecting ordinal position in the sequence. We test the generalizability of this so-called primacy gradient across motor sequences and memorized auditory sequences. Using a multivariate decoding approach, Experiment 1 replicated the presence of a Competitive Queuing primacy gradient in magnetoencephalography (MEG) data of young healthy adults (n = 23) when they prepared a sequence of finger movements from memory. Importantly, we observed a similar primacy gradient when participants anticipated a sequence of tones they had learned before, in the absence of any movement. In Experiment 2 (n = 23; naïve cohort), we rule out the possibility that this primacy gradient in auditory memory is explained by any learnt association between tones and movements, or by MEG signal fluctuations that are unrelated to discrete sequential events. In sum, we find a similar neural signature of serial order coding when humans prepare a sequence of movements, and when they anticipate a sequence of sounds. This lends support to the generalizability of Competitive Queuing.

## 2 Introduction

Across species, behavior often consists of sequences of events that follow a certain order, from self-grooming to communicating with others. Humans learn exceptionally complex behavioral sequences that structure both our actions and our perceptual experience, such as in language or music. Yet it remains unknown whether sequences of events we perceive, and sequences of our own actions, are encoded similarly in the brain.

Behavioral and theoretical work suggests that the brain may represent serial order through a domain-general neural mechanism consistent with the computational principle of Competitive Queuing (CQ; introduced by Grossberg [1]; term coined by Houghton [2]; for a review see Hurlstone et al. [3]). In its basic form, CQ involves a parallel planning layer and a competitive choice layer. In the parallel planning layer, all sequence items are represented simultaneously in separate nodes, with activation strength decreasing according to their serial position. This simultaneous graded activation pattern establishes a queue of upcoming items. The choice layer turns this pattern into a sequence of events through competing inhibitory interactions between sequence items, allowing only the item with the currently highest activation strength to be selected at a time. Then, self-inhibitory feedback suppresses the selected item so the next strongest item in the queue can follow.

CQ models can explain benchmarks of human behavior observed in serial order working memory tasks, such as primacy effects and transposition errors, among others [3,4]. Such behavioral phenomena exist in the verbal, visual, and spatial domains (comprehensive review in Oberauer et al. [4]), as well as the motor domain [5,6]. This has motivated the idea that CQ may be, at least partly, domain-general [3]. Indeed, a wide range of serial behaviors and mental faculties have been modeled with the CQ principle. The range of serial behaviors examined includes word list memorization [7], daily actions [8], perception of musical melodies [9], motor sequences [10], and sequences with a temporal component, i.e. rhythmic pattern [11,12].

Surprisingly, while theoretical work on CQ has focused on short-term memory across multiple domains, direct neurophysiological evidence for CQ stems only from the motor domain. Averbeck et al. [13] provided evidence for graded parallel representations of forthcoming actions in monkeys preparing to draw geometrical shapes (e.g., a square or a triangle). The authors showed that neurons in prefrontal cortex represented several upcoming movement segments simultaneously, at a time when the monkey already knew which shape to draw but had not yet initiated the drawing movement. Importantly, the strength of each representation scaled with the position of its associated movement segment in the upcoming sequence, as predicted by CQ. Using magnetoencephalography (MEG), Kornysheva et al. [14] reported a similar pattern when humans prepared previously learned sequences of finger movements. Evidence for a role of CQ in the preparation of motor sequences also comes from behavior [15–17], from motor evoked potentials [18,19], and from the lateralized readiness potential [20,21]. Direct neurophysiological evidence for CQ in non-motor domains is, to our knowledge, pending. This limits the generalizability of CQ as a principle underlying neural coding of serial order across domains.

Here, we provide evidence that CQ extends to the neural coding of serial order in a perceptual task that relies on serial order memory. In Experiment 1, we replicate previous findings of a parallel, graded representation of serial order when humans prepare to execute a sequence of finger movements. Importantly, we observed a comparable parallel gradient during cued anticipation of sequences of sounds. In Experiment 2, we rule out the possibility that this parallel gradient during anticipation of sounds arises from any auditory-motor associations in Experiment 1 [22,23], or from features of the task design unrelated to serial information. Our findings establish CQ as a general principle underlying neural codes of serial order.

## 3 Results

We designed an MEG experiment in which participants prepared a known sequence of finger movements, or anticipated a learnt sequence of tones. Prior to the start of the experiment, participants learned to uniquely associate each of two visual cues (abstract fractals) with a distinct sequence of five keypresses, and each keypress with a distinct tone (notes C to G, mapped in ascending order onto the five fingers of the left hand, starting with the little finger). The experiment consisted of two tasks, a motor sequence task and a tone sequence task. In both tasks, each trial started with the presentation of one of the visual cues. In the motor sequence task (**Fig 1A**), the cue informed participants which sequence of keypresses to prepare for execution. Once the visual cue disappeared (after 1.8 to 2.2 seconds), participants had to execute that sequence from memory at a regular pace of 2 Hz. There were two versions of this task, one in which each keypress generated its associated tone, and one in which keypresses triggered no tones (silent production). Results were similar for the two versions. We therefore focus on the silent production version (see supplementary material for results from the version that included tones). In the tone sequence task (**Fig 1B**), the visual cue informed participants which tone sequence they would hear once the cue disappeared (at a pace of 2 Hz). Rare deviant sequences (mismatches between the cued and presented tone sequence) ensured that participants actively anticipated the cued tone sequence during the preparation phase and remained engaged throughout the trial. In both the motor sequence task and the tone sequence task, visual cues alternated every three trials. Tasks alternated across blocks of 24 trials each.

**Fig 1.**
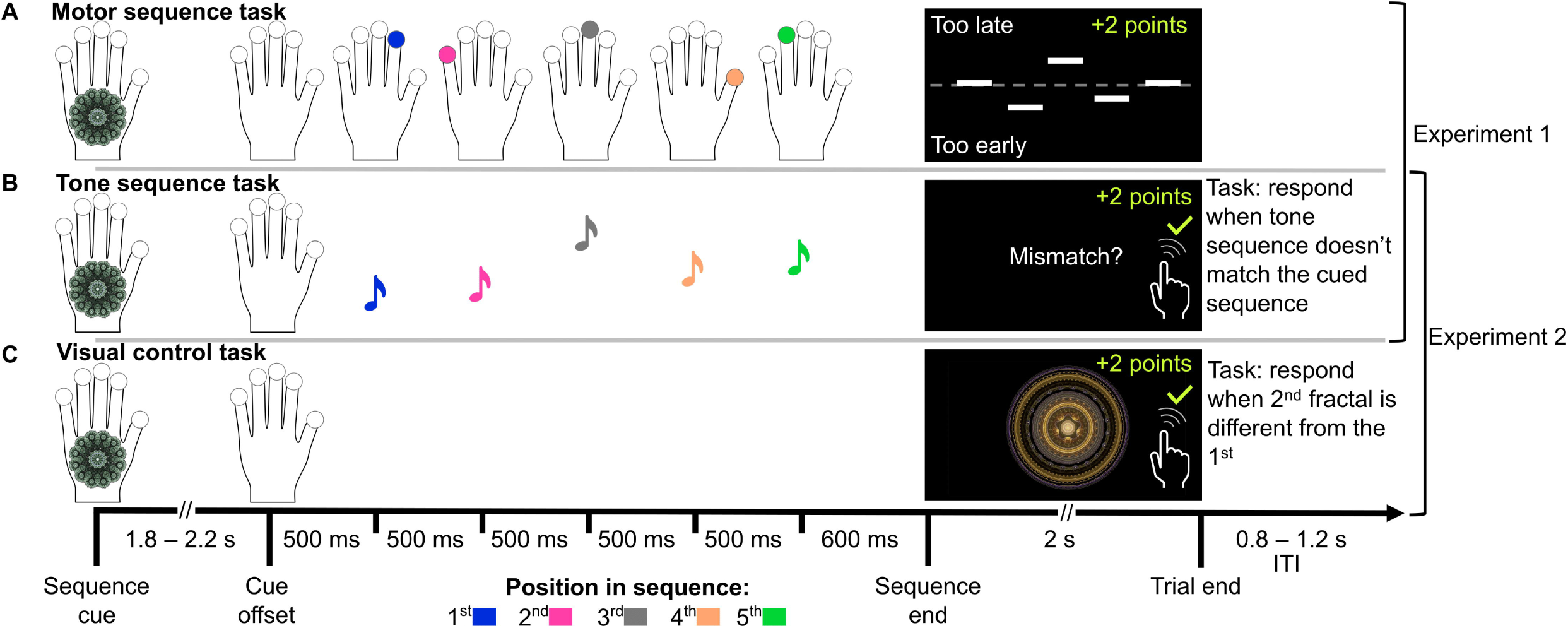
Task structure. In Experiment 1, participants completed a motor sequence task and a tone sequence task while we recorded MEG. In each trial of the motor sequence task (**A**), participants saw a visual cue, and prepared to execute a sequence of five keypresses which they had learnt to associate with that cue. When the visual cue disappeared (after 1.8-2.2 s), participants executed the keypress sequence at a pace of 2 Hz, starting 500 ms after cue offset. At the end of each trial, participants received visual feedback indicating whether each keypress was correct, and how accurate its timing was. The tone sequence task (**B**) used the same two visual cues as the motor sequence task. Here, the visual cue at trial start informed participants which sequence of five tones they would hear, at a pace of 2 Hz, when the cue disappeared (starting 500 ms after cue offset). While listening to the tone sequence, participants had to refrain from any movement, and detect rare mismatches between the cued and presented tone sequence. This ensured that participants retrieved the tone sequence associated with the visual cue from memory. Experiment 1 established an intuitive correspondence between the order of keypresses associated with a given cue, and the order of tones associated with that same cue, because participants learnt before the experiment to associate each keypress with a unique tone (notes C to G, mapped in ascending order onto the five fingers of the left hand, starting with the little finger). Panel B shows an example of the rare mismatches between the cued and presented tone sequence (the vertical position of the notes illustrate the pitch of the corresponding tone, i.e., the orange note, associated with the thumb, should be highest). Experiment 2 did not include any motor sequence task, and thus eliminated any potential alternative explanation that arose from this link between finger movements and tone sequences. In Experiment 2, participants memorized the tone sequences and the cue-sequence association independently of any finger movement, through pure listening, and then performed the tone sequence task as in Experiment 1. **C:** To rule out any influence of systematic MEG signal fluctuations across the course of a trial that are unrelated to discrete sequential events, participants additionally performed a visual control task, in which the visual cue was followed by no sequential keypresses or tone sequence. A visual mismatch task, in which participants compared the initial visual cue at trial start with a fractal shown at trial end, ensured participants’ task engagement (mismatch trial illustrated in panel C). Participants performed the visual control task first, while they were still naïve to any association between the visual cue and any sequential information. For all tasks, we provided performance feedback after every trial. Fingers and tones are color-coded for rank order for illustration purposes only.

### 3.1 Experiment 1

#### 3.1.1 Behavior

In the motor sequence task, participants produced correct keypress sequences in 94.4% of trials (mean across participants; SD = 5.4%, range: 75 to 100%). Keypress speed was faster than instructed, with an average inter-press interval of 475 ms (median across keypresses, mean across participants; SD = 22 ms, range: 428 to 510 ms; **Fig 2A**). Inter-press intervals varied, on average, by 85 ms (IQR across keypresses, mean across participants; SD = 28 ms, range: 42 to 152 ms). In the tone sequence task, all participants successfully detected mismatches between cued and presented tone sequences (mean d’ = 3.112, SD = 1.048, range: 1.569 to 5.399; **Fig 2B, left**).

**Fig 2.**
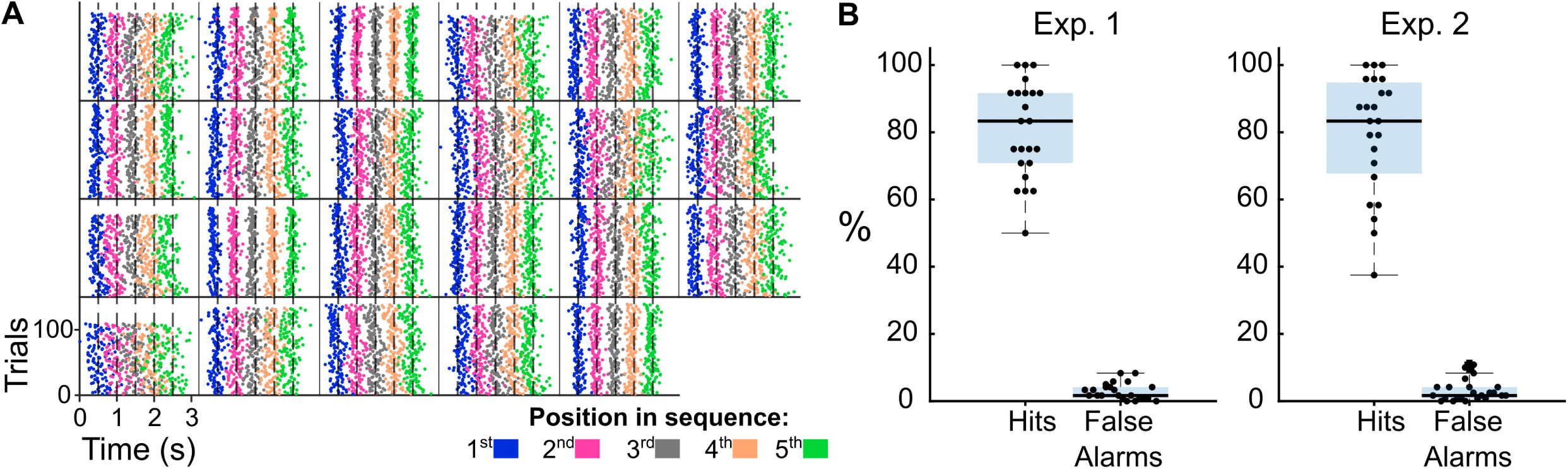
Behavioral performance. **A:** Individual participant plots (n=23) from Experiment 1, showing the produced keypresses in the motor sequence task. Each plot shows for each trial (y-axis) the time of each press (x-axis) relative to the fractal offset, with color coding for the position of each press in the sequence. Dashed vertical lines indicate the target time points at the target keypress rate of 2 Hz. **B:** Boxplots for performance of detecting mismatches during the tone sequence task in Experiment 1 (**left**) and Experiment 2 (**right**), shown as Hit Rates (percentage of deviant trials that each participant successfully reported as deviant) and False Alarm Rates (percentage of non-deviant trials that each participant wrongfully reported as deviant). From these rates we compute the sensitivity index d’ (see Methods).

#### 3.1.2 MEG

We tested whether anticipation of an upcoming sequence of tones evokes simultaneous mental representations of those tones, graded by their expected position in the sequence. Such a parallel, anticipatory gradient would point to a similar principle underlying the neural representation of serial order during anticipation of sequential sensory input as previously described during preparation of sequential movements [13,14]. To identify parallel gradients in the tone sequence task, and in the motor sequence task, we employed a multivariate decoding approach. This approach used the positions of consecutive items in a sequence (tones, or movements) as training labels (1^st^ to 5^th^ item), and the 102 magnetometer channels as features. For each channel, we computed the mean MEG signal across the 100 ms before each item onset (**Fig 3**; see also [14]).

**Fig 3.**
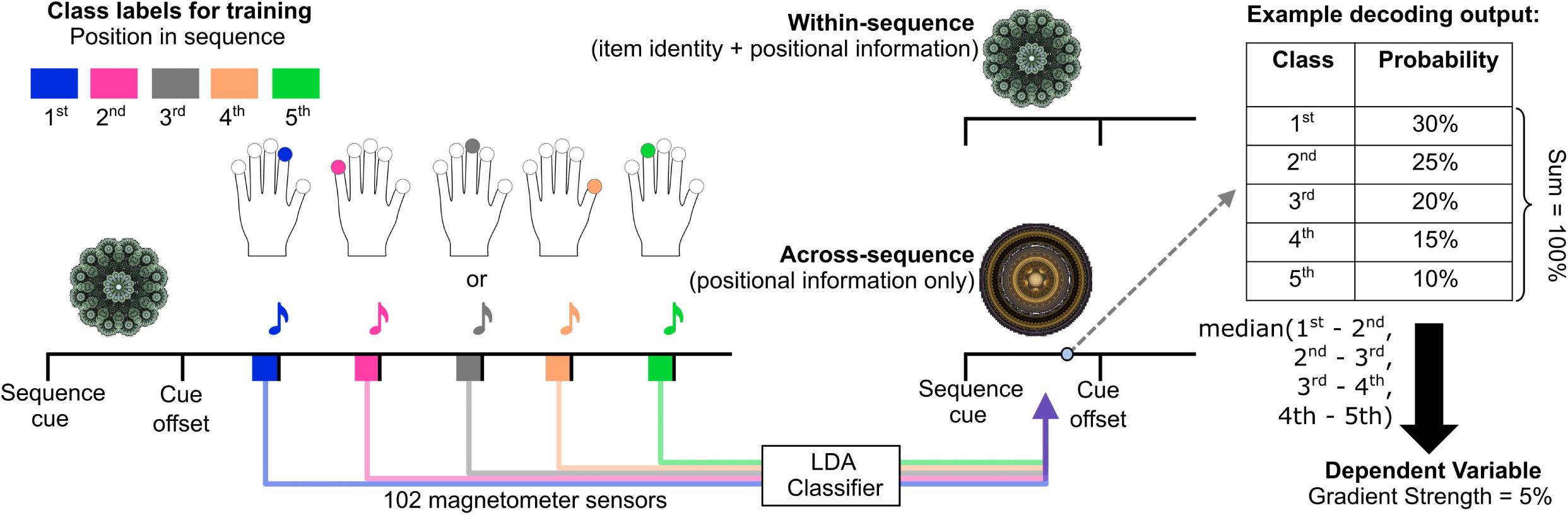
MEG data analysis. We trained an LDA classifier to discriminate items of a sequence using as training labels the item’s position in the sequence, and as features the 102 magnetometer sensors, averaging the signal over the last 100 ms before keypress or tone onset per channel. Decoding was performed separately for the motor sequence task and the tone sequence task. We decoded every 10-ms time bin of each trial, obtaining as an output from the classifier the probability that each time point is classified as each of the 5 classes (called pattern probabilities throughout the manuscript). Our ‘within-sequence’ decoding approach trained and tested classifiers on trials that shared the same sequence, using 5-fold cross-validation. Our ‘across-sequence’ decoding approach trained classifiers on all trials of one sequence, and then tested all trials of the other sequence to isolate positional information from information about specific movement effectors, or specific tones. Here, we illustrate as an example the pattern probabilities for a single time point of the preparation window. In this example, the probabilities show a gradient that aligns with previous evidence of the Competitive Queuing mechanism [13,14], i.e., each position’s probability of being decoded reflects its rank in the sequence. Our final metric, Gradient Strength, is the median difference of successive pairs of probabilities (i.e., 1^st^ minus 2^nd^, 2^nd^ minus 3^rd^, etc., see [14]), summarized across trials, time points of the time window of interest (last second before cue offset), and sequences (and repetitions for the ‘within’ cross-validated decoding). Another quantification for a second time window of interest, the last 800 ms before cue onset, i.e., the ITI, was used as a baseline to compute the baseline-corrected Gradient Strength.

First, we confirmed that our LDA classifier could discriminate the position of a tone, or a movement, either within the same sequence (**Fig 4A-F**), or across different sequences of tones or movements, respectively (**Fig 4G-L**). While within-sequence decoding does not separate information about position from information about specific movements, or specific tones, training a classifier on one sequence and testing on another sequence, which has a different order of movements, or tones, isolates positional information [14].

**Fig 4.**
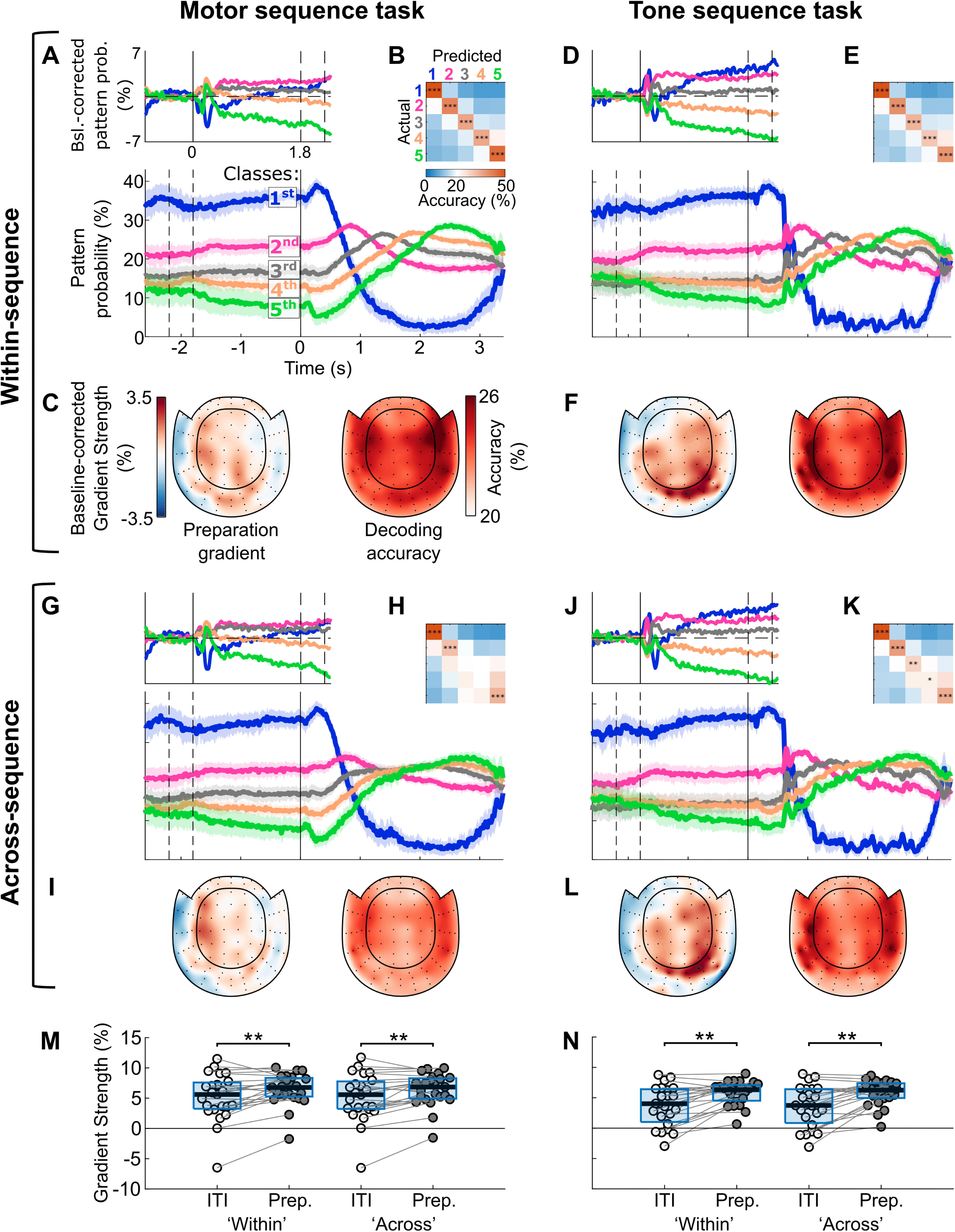
A parallel gradient represents serial order in the motor sequence task (left column) and the tone sequence task (right column) in Experiment 1. **A-F:** within-sequence decoding (sequence A=>A, and B=>B), 5-fold cross-validated. **A, lower panel** shows grand-average time-courses of pattern probabilities for each of the five classes, with color coding for class, i.e., position in the sequence. The time axis is time-locked to cue offset, which signaled the start of the response window in the motor sequence task. The dashed lines at t = -1.8 and -2.2 s indicate the range of the possible cue onsets (fractal duration was jittered). Shadings represent 95% confidence intervals. **A, upper panel:** Difference in pattern probabilities from baseline (-0.8 to 0 s relative to cue onset), time-locked to cue onset. The dashed lines at t = 1.8 and 2.2 s indicate the range of the possible cue offsets. Time-courses in panel A, upper, are smoothed with a Gaussian window spanning five bins (=50 ms). **B,** grand-average confusion matrix of the classifier’s cross-validated decoding performance in discriminating items of the same sequence. Elements of the diagonal are tested against chance level (20%), and *p*-values are Bonferroni-Holm corrected for five comparisons. **C,** searchlight results, indicating which sensors show the highest baseline-corrected Gradient Strength (left), and which sensors show the highest decoding accuracy when classifying sequence items during the response window (right). **D-F,** same as A-C, but for the tone sequence task ‘within’ decoding. **G-L:** same as A-F, but for across-sequence decoding (sequence A=>B, and B=>A). **M-N,** Quantification of the degree to which pattern probabilities are graded by rank in the motor sequence task (M) and tone sequence task (N). The y-axis shows Gradient Strength, i.e., the median difference of pattern probabilities between successive pairs of positions per participant, summarized across trials, sequences (and repetitions for ‘within’ decoding), and time bins from two time windows of interest: the baseline (‘ITI’) and the last second before cue offset (‘Prep.’). Gradient Strength was significantly above zero even during the ITI, possibly due to the predictable switching of sequences in our paradigm which may have allowed participants to predict the upcoming sequence well before the cue onset. The box edges define the upper and lower quartiles. The horizontal black lines indicate the group median. We compared the Gradient Strength between the baseline and the last second before cue offset, for both tasks and decoders, with paired-samples t-tests (*p*-values Bonferroni-Holm corrected for four comparisons). * *p* < 0.05, ** *p* < 0.01, *** *p* < 0.001, Bonferroni-Holm corrected.

In the motor sequence task, mean decoding accuracy across participants was 44.78% ± 6.77% for within-sequence classification, and 34.02% ± 5.15% for across-sequences classification. In the tone sequence task, the classifier achieved an accuracy of 40.95% ± 7.74% within-sequences, and 37.32% ± 5.17% across-sequences. When testing decoding accuracies for each sequence position separately against chance level (20%) at the group level, we found above chance-level within-sequence classification for all sequence positions, both in the motor sequence task (**Fig 4B**), and the tone sequence task (**Fig 4E**; motor sequence task: *p_corr._* < 0.001 for all positions; tone sequence task: *p_corr._* < 0.001 for the first, second, and fifth position, *p_corr._* = 0.019 for the third position, and *p_corr._* = 0.022 for the fourth position; Bonferroni-Holm corrected for five comparisons, separately for each task). Classification across-sequences (**Fig 4H, K**), deteriorated for the third and fourth serial positions in motor sequences (*p_corr._* < 0.001 for the first, second, and fifth position, *p_corr._* = 0.891 and *p_corr._* = 0.590 for the third and fourth position, respectively; Bonferroni-Holm corrected for five comparisons), but not in tone sequences (*p_corr._* < 0.001 for the first, second, and fifth position, *p_corr._* = 0.001 and *p_corr._* = 0.035 for the third and fourth positions, respectively; Bonferroni-Holm corrected for five comparisons).

Having confirmed that our classifiers could discriminate elements of tone sequences as well as movement sequences during listening and production, respectively, we tested for a mental representation of sequence elements during preparation, i.e., in response to the cue, before any motor output or auditory input. For each 10-ms time bin of the preparation time window, we obtained five probability values. Each probability indicated how likely it was that the MEG data in that time bin represented the corresponding sequence item.

For the motor sequence task, this analysis confirmed that preparation entailed a parallel gradient, i.e., simultaneous representations of several movements, and scaling of the probability of representation with the position of a movement in the upcoming sequence (**Fig 4A,G**). To quantify the degree to which preparatory representations followed a parallel gradient, we computed Gradient Strength as the median difference between probabilities of successive sequence elements (1^st^ minus 2^nd^, 2^nd^ minus 3^rd^, etc. [14]), from two time windows of interest, i.e. the last second before cue offset and the baseline. Gradient Strength was significantly above zero for both time windows, and for both decoders (within-sequence and across-sequence decoding; **Table 1)**. Given the predictable order of the two sequences during the experiment (sequences A and B alternated every three trials), participants could form expectations about the upcoming cue, and its associated sequence, even before the presentation of the cue, which could explain why Gradient Strength was significantly above zero even before cue onset. Importantly, Gradient Strength significantly increased after cue onset, compared to baseline, indicating that participants started fully preparing for the upcoming sequence only after the visual cue was presented (2x2x2 repeated-measures ANOVA with the within-subject factors Decoder, Task, and Phase: main effect of Phase, *F*_(1,22)_ = 19.247, *p* < .001, 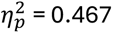; **Table 1** and **Fig 4M**). There was no decrease in Gradient Strength when decoding across sequences, compared to decoding within sequences (no significant main effect of Decoder, or any interaction involving Decoder; *F* < 0.4, *p* > 0.5), indicating that the parallel gradient was largely independent of information about the specific movements in a sequence, and instead reflected positional information. Together, our findings in the motor sequence task replicate previous findings that preparation for motor sequences evokes a parallel gradient, in line with Competitive Queuing [13,14]. Results from the version of the motor sequence task in which keypresses triggered tones were qualitatively similar, and are reported in the Supporting Information (**Fig 1 in S1 Supporting Information**).

**Table 1.**
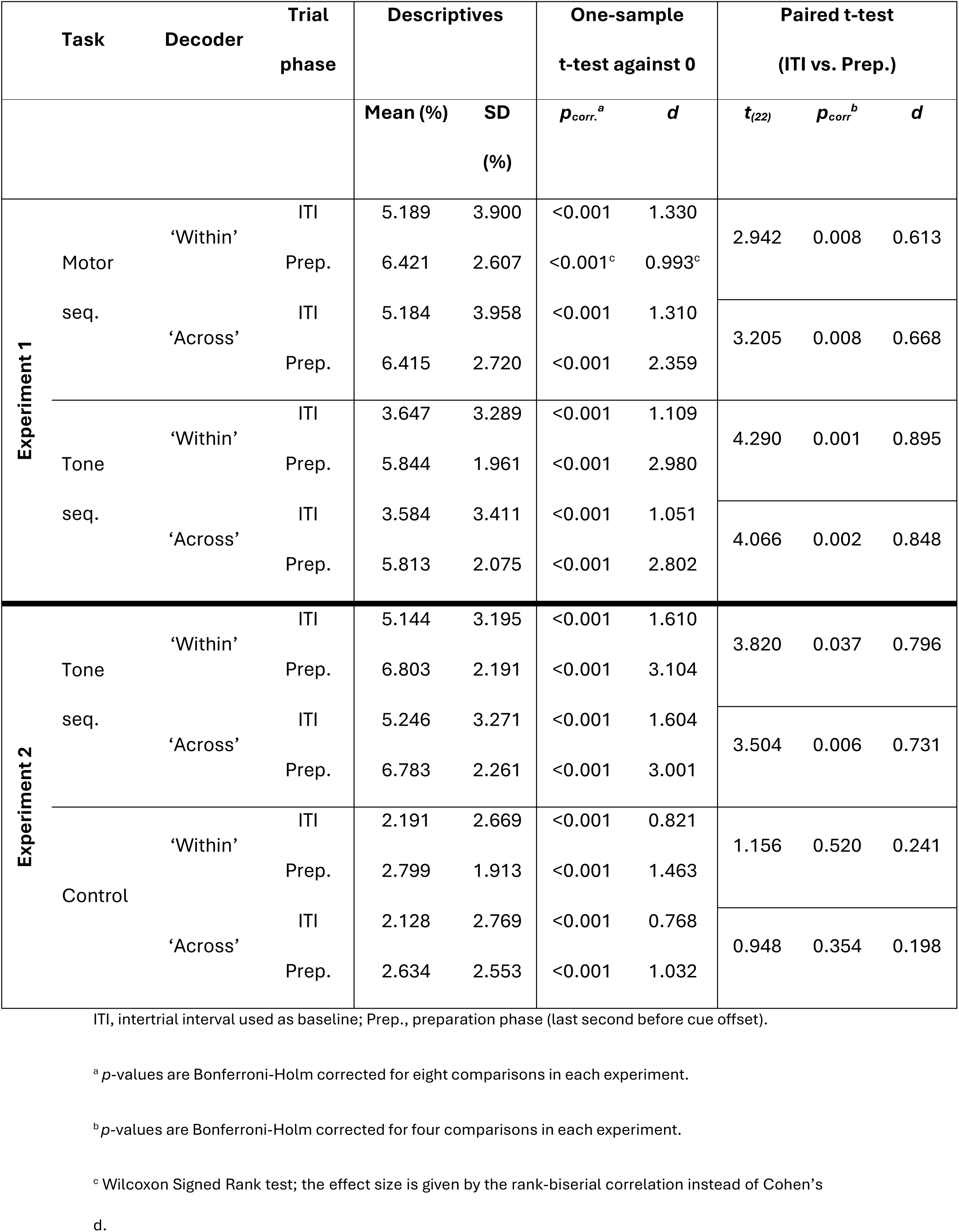
Summary of statistics on Gradient Strength from both experiments.

Importantly, we found a similar parallel gradient of positional information in the tone sequence task (**Fig 4D,J**). Gradient Strength was again significantly above zero in the last second before cue offset, as well as in the baseline, and significantly increased from baseline to the last second before cue offset (main effect of Phase reported in the previous paragraph; **Table 1** and **Fig 4N**). Gradient Strength reflected information about the ordinal positions of tones, with no indication of tone-specific coding (no main effect of Decoder, or any interaction involving Decoder, as reported in the previous paragraph). The main effect of Task, and the Task x Phase interaction were not significant (*F* > 3.7; *p* > 0.065). A control analysis, which excluded trials with above-threshold electromyography (EMG) activity in left hand and forearm muscles, ruled out muscle activity as a major driver of Gradient Strength (**Fig 2 in S1 Supporting Information**). In sum, results supported our hypothesis that a parallel gradient is a domain-general principle of serial order representation during preparation for sequences of movements, and sequences of sensory input.

A correlation analysis of individual differences across the two tasks provided further support for this domain-generality. Participants who showed a pronounced increase in Gradient Strength from the ITI to the last second before cue offset in one task also had a pronounced increase in the other task, independently of the decoding approach (‘within’: *r* = 0.423, *p* = 0.044; ‘across’: *r* = 0.424, *p* = 0.044; **Fig 5, top row**).

**Fig 5.**
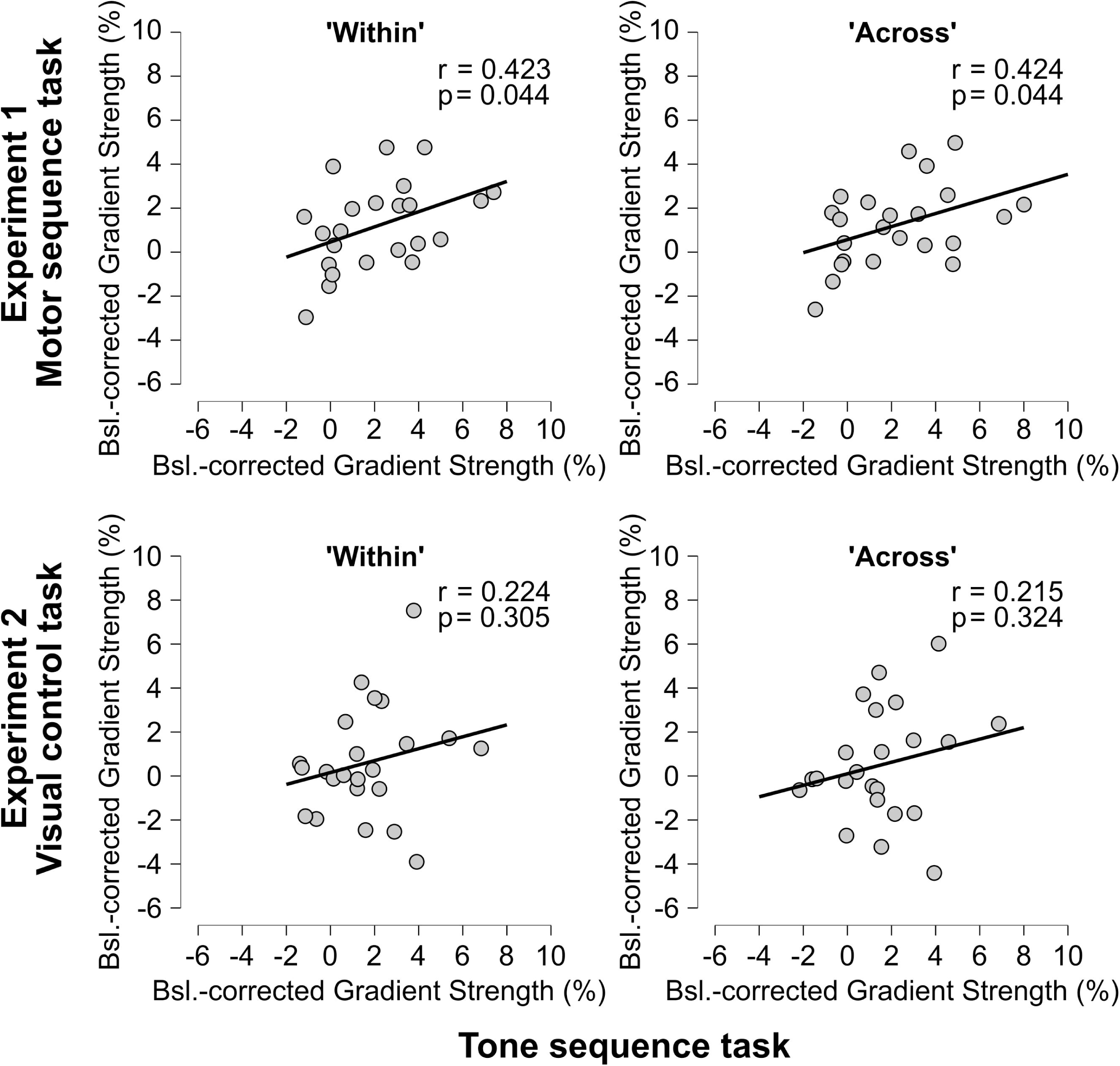
Correlation analysis of serial order representations between tasks. **Top row**: Correlation between tone sequence and motor sequence task in Experiment 1. Baseline-corrected Gradient Strength in one task is positively associated with baseline-corrected Gradient Strength in the other task, hinting at a domain-general mechanism for representing serial order during the preparation of sequences. **Left:** within-sequence decoding; **Right:** across-sequence decoding. **Bottom row**: Correlation between tone sequence and visual control tasks in Experiment 2. The correlation is not significant for either ‘within’ (**Left**) or ‘across’ (**Right**) decoding. This analysis further corroborates that it is the presence of serial order information that drives the Gradient’s Strength increase from baseline.

To assess possible differences in the anatomical basis of the gradient across domains, we compared the searchlight topographies of baseline-corrected Gradient Strength (**Fig 4C,F,I,L, left topographies**) between the motor production task and the tone listening task, using a within-subject, two-tailed t-test in a Montecarlo-based cluster-based permutation test. This analysis revealed a significant cluster (‘within’: *p* = 0.008; ‘across’: *p* = 0.005) involving right lateral frontal and temporal sensors, where the baseline-corrected Gradient Strength was greater for the tone sequence task than for the motor sequence task (**Fig 6, top row**). While we do not make precise anatomical claims, the mere existence of topographical differences suggests partly domain-specific components in the mechanism of serial order memory.

**Fig 6.**
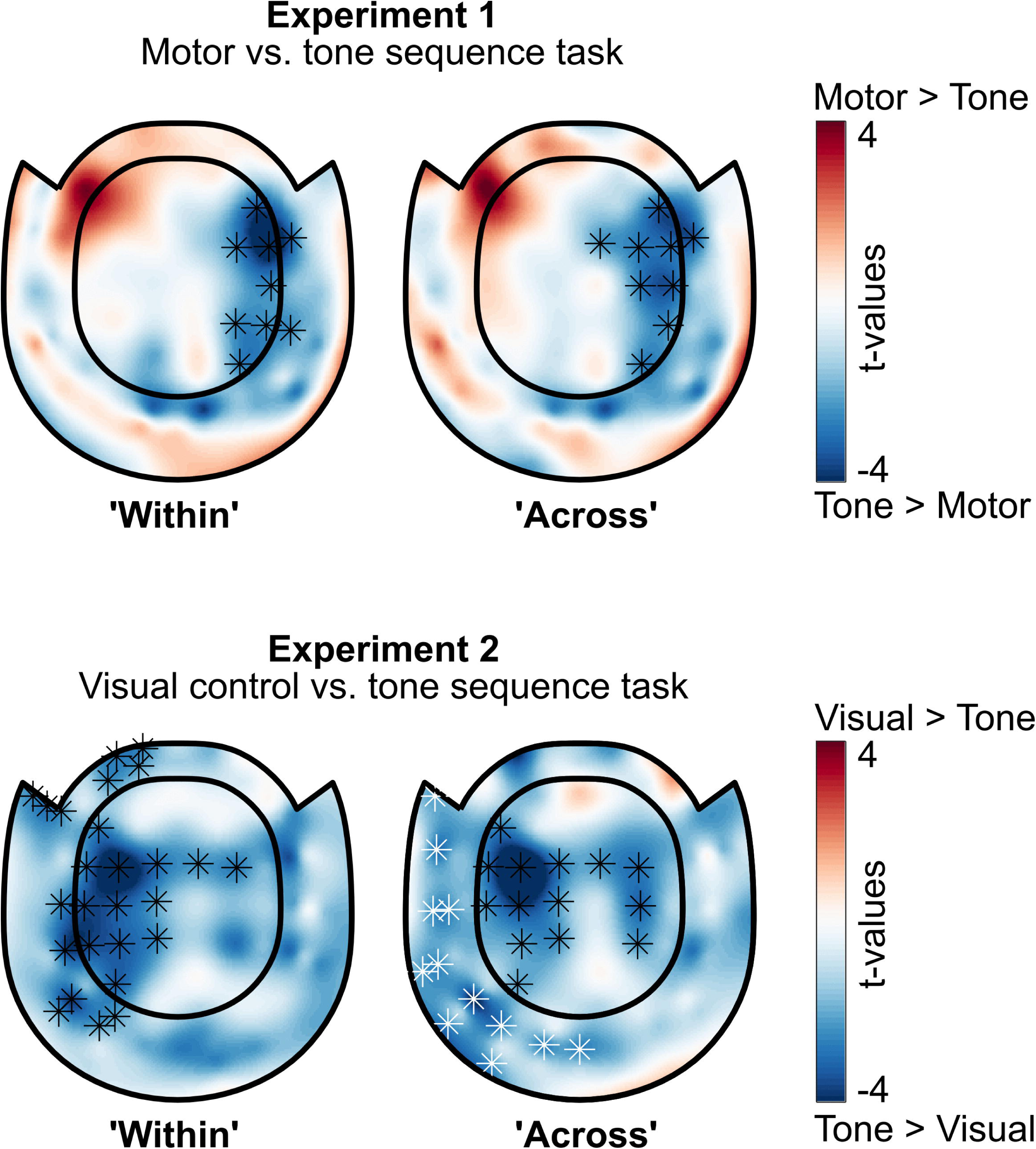
Differences in the topography of baseline-corrected Gradient Strength between the tasks. **Top row**: comparison between motor sequence and tone sequence tasks in Experiment 1. A significant cluster over the right hemisphere shows an increased change from baseline in the tone sequence task, compared to the motor sequence task. **Bottom row**: comparison between visual control and tone sequence tasks in Experiment 2. A significant cluster over the left hemisphere shows increased change from baseline in the tone sequence task, compared to the visual control task, when decoding within-sequence. Across-sequences, two clusters over the left hemisphere (separated by color: black and white markers) show increased change from baseline in the tone sequence task, compared to the visual control task.

### 3.2 Experiment 2

Experiment 2 aimed to replicate the finding of a Competitive Queuing gradient when participants anticipate a sequence of tones, and to rule out two potential sources of ambiguity in the findings. These were the possibility that motor representations emerged during the anticipation of a tone sequence due to association of motor responses with tones in Experiment 1, and the possibility that classifiers were influenced by systematic MEG signal fluctuations not directly related to the mental representations of the tone sequence. Thus, we tested a naïve cohort in the same tone sequence task as in Experiment 1 while avoiding any mapping of tones to movements. We also included a control task that was devoid of any sequence-related information, which participants carried out at the start of the session before we established any cue-sequence associations. The control task was a delayed match-to-sample task. Participants saw a visual cue at trial start (one of the two fractals used in the tone sequence task), then waited for 3.1 seconds after cue offset (the time window during which tones were presented during the tone sequence task) before viewing a second fractal, which they had to judge as identical, or different, compared to the first fractal. Thus, trial events and event timings up to the second fractal were identical to the tone sequence task, except that the control task did not involve any discrete sequential information during the 3.1-second waiting period.

#### 3.2.1 Behavior

In Experiment 2, the tone sequence task was identical to Experiment 1, with the exception that participants learned the tone sequences without association to any motor sequence. Participants successfully learned the mapping of visual cues to tone sequences, evidenced by high accuracy in detecting tone sequence mismatches (average d’ = 3.118, SD = 1.146, range: 1.235 to 5.399; **Fig 2B, right**). In the delayed match-to-sample control task, participants detected mismatches between the first and second fractal with high sensitivity (average d’ = 4.141, SD = 0.987, range: 2.559 to 5.399).

#### 3.2.2 MEG

Similar to Experiment 1, we evaluated the classifier’s performance when predicting serial positions either within-sequence or across-sequences. Classification of serial positions of tones in the tone sequence task was successful, with an average decoding accuracy of 38.33% ± 4.78% within-sequence, and 37.57% ± 4.94% across-sequences. For the control task, we defined mock events at the five time points relative to cue offset at which tones were presented in the tone sequence task. In the control task, these time points were not associated with any discrete event. Decoding accuracies for these mock events were, on average, 27.81% ± 4.20% for both within- and across-sequence decoding. Testing each item’s decoding accuracy against chance level (20%) revealed successful classification for all positions in the tone sequence task within-sequence (**Fig 7B**; *p_corr._* < 0.001 for all positions, Bonferroni-Holm corrected for five comparisons), and above chance accuracy for all positions except the fourth across-sequences (**Fig 7H**; *p_corr._* < 0.001 for the first, second, and fifth position, *p_corr._* = 0.001 for the third position, and *p_corr._* = 0.149 for the fourth position, Bonferroni-Holm corrected for five comparisons). In the control task, decoding accuracy was above chance for three of the five positions, with no difference between within- and across-sequence decoding (within: **Fig 7E**, *p_corr._* < 0.001 for the first and fifth positions, *p_corr._* = 0.003, *p_corr._* = 0.982, *p_corr._* = 0.618, for the second, third, and fourth positions respectively; across: **Fig 7K**, *p_corr._* < 0.001 for the first and fifth positions, *p_corr._* = 0.006, *p_corr._* = 0.939, *p_corr._* = 0.863, for the second, third, and fourth positions respectively; *p*-values Bonferroni-Holm corrected for five comparisons, for each decoding approach separately).

**Fig 7.**
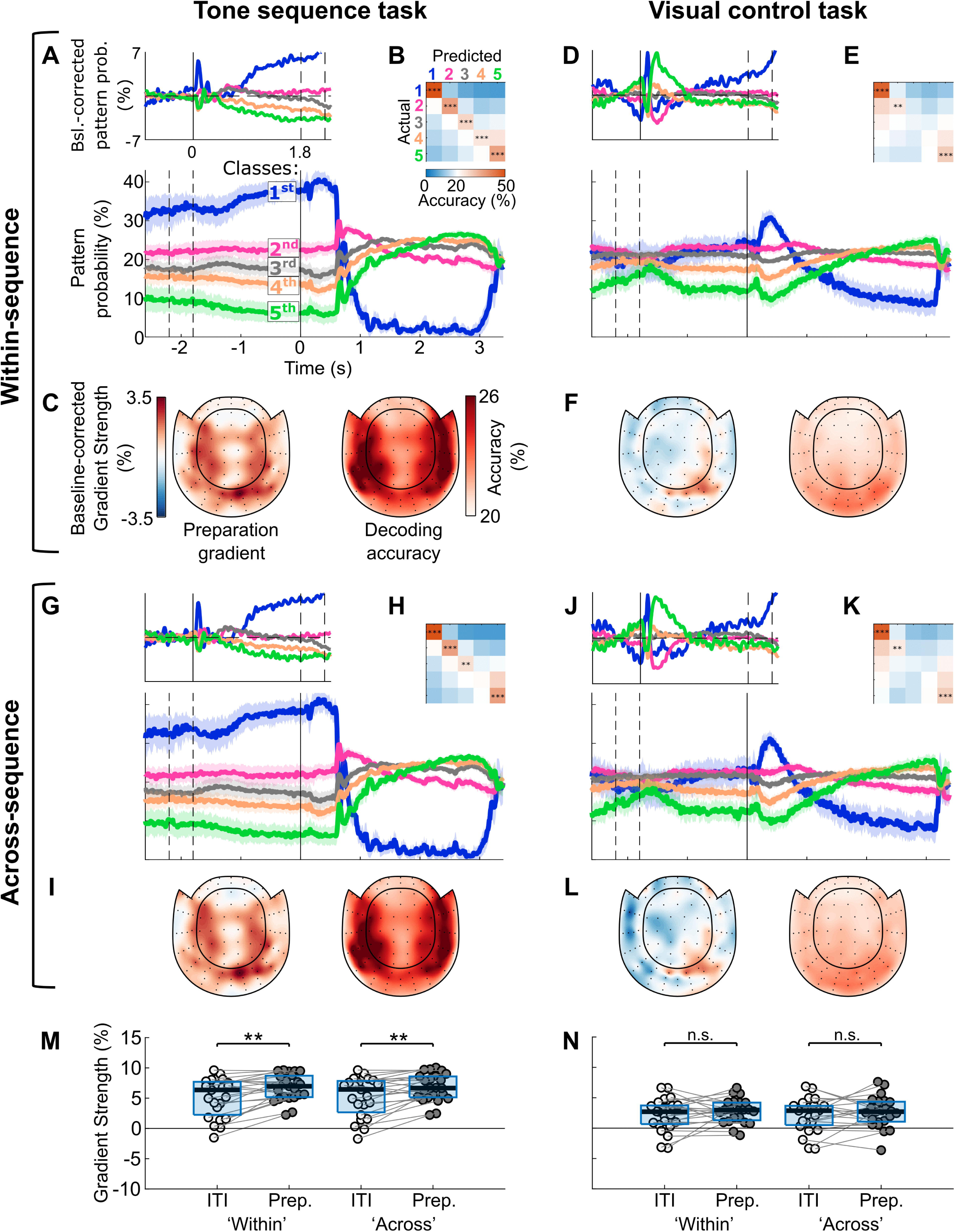
The parallel gradient in the tone sequence task in Experiment 2 persists in the absence of any mapping of tones to movements (left column), and it is reduced in the absence of any planning task (visual control; right column). **A-F,** within-sequence decoding (sequence/cue A=>A, and B=>B), 5-fold cross-validated. **A, lower panel** shows the grand-average time-course of pattern probabilities for each of the five classes, with color coding for class, i.e., position in the sequence. The time axis is time-locked to cue offset, which signaled the start of the presentation window in the tone sequence task. The dashed lines at t = -1.8 and -2.2 s indicate the range of the possible cue onsets (fractal duration was jittered). Shadings represent 95% confidence intervals. **A, upper panel,** difference in pattern probabilities from baseline (-0.8 to 0 s relative to cue onset), time-locked to cue onset. The dashed lines at t = 1.8 and 2.2 s indicate the range of the possible cue offsets. Time-courses in panel A, upper, are smoothed with a Gaussian window spanning five bins (=50 ms). **B,** group-average confusion matrix of the classifier’s cross-validated decoding performance to discriminate items of the same sequence. Elements of the diagonal are tested against chance level (20%), and *p*-values are Bonferroni-Holm corrected for five comparisons. **C,** searchlight results, indicating which sensors show the highest baseline-corrected Gradient Strength (left), and which sensors show the highest decoding accuracy when classifying sequence items during the presentation window (right). **D-F,** same as A-C, but for the visual control task ‘within’ decoding. **G-L,** same as A-F, but for across-sequence decoding (sequence/cue A=>B, and B=>A). **M-N,** Quantification of the degree to which pattern probabilities are graded by rank in the tone sequence task (M) and control task (N). The y-axis shows Gradient Strength for two time windows of interest: the baseline (‘ITI’) and the last second before cue offset (‘Prep.’). The box edges define the upper and lower quartiles. The horizontal black lines indicate the group median. We compared Gradient Strength between the ITI and the last second before cue offset, for both tasks and decoders, with paired-samples t-tests (*p*-values Bonferroni-Holm corrected for four comparisons). n.s. = not significant, * *p* < 0.05, ** *p* < 0.01, *** *p* < 0.001, Bonferroni-Holm corrected.

Next, as in Experiment 1, we decoded all time bins of the trial epoch (**Fig 7A,D,G,J**) and quantified the gradient as Gradient Strength for all decoders, tasks and time windows of interest. Gradient Strength was significant in the tone sequence task, but also in the control task, for both the baseline and the last second before cue offset (all Gradient Strengths were significantly above zero; **Table 1**). However, a 2x2x2 repeated ANOVA (within factors: Decoder, Task, Phase), and planned comparisons for Phase, revealed a stronger gradient in the tone sequence task, compared to the control task, and a significant increase in Gradient Strength from baseline to the last second before cue offset only in the tone sequence task. Specifically, we found a main effect of Task (*F*_(1,22)_ = 35.302, *p* < .001, 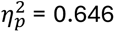), showing that Gradient Strength was higher in the tone sequence task than in the control task. We also found a main effect of Phase (*F*_(1,22)_ = 8.244, *p* < .009, 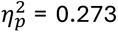), showing that Gradient Strength increased from the baseline to the last second before cue offset. However, a trend of Task x Phase interaction (*F*_(1,22)_ = 3.026, *p* < .096, 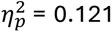), as well as planned comparisons for Phase in both tasks (**Table 1**, and **Fig 7M-N**), revealed that the main effect of Phase was driven by changes from baseline in the tone sequence task alone, while there was no significant change from baseline in the control task. Lastly, the correlation of baseline-corrected Gradient Strength between tasks was not significant (‘within’: Pearson’s *r* = 0.224, *p* = 0.305; ‘across’: *r* = 0.215, *p* = 0.324; **Fig 5, bottom row**).

As in Experiment 1, we compared baseline-corrected Gradient Strength topographies across tasks (**Fig 6, bottom row**). While in Experiment 1 we compared two tasks that both require serial order processing, thus highlighting potential domain-specific topographical differences, comparing the two tasks in Experiment 2 highlights the topographical differences of presence vs. absence of serial order processing. The analysis revealed a significant - mostly right-lateralized - cluster of sensors that showed higher change from baseline for the tone sequence task than the visual task when decoding within-sequence (*p* = 0.001), and two clusters when decoding across-sequence, with one cluster including mostly right temporal and temporooccipital sensors (*p* = 0.008), and the other cluster including left and right central sensors (*p* = 0.004).

Taken together, these results confirm a parallel gradient representing serial order during anticipation of the tone sequence. Even though the control task captured some sequence-unrelated MEG activity that gave rise to a pattern resembling a gradient, this gradient was significantly weaker than in the tone sequence task and did not show the significant increase upon cue presentation that characterized the gradient in the tone sequence task. We therefore conclude that sequence-unrelated activity cannot fully explain the gradient in the tone sequence task.

## 4 Discussion

Based on the hypothesis that Competitive Queuing is a domain-general principle of serial order coding in the brain, we tested for neurophysiological evidence of CQ in a motor sequence task and an auditory sequence memory task. In both tasks, we found parallel representations of several items of an upcoming sequence while participants prepared for that sequence, i.e., before the first movement in the motor sequence task, and the first tone in the auditory memory task. The representational strength of each item scaled with its serial position in the upcoming sequence, in line with the primacy gradient predicted by CQ models [3]. Our control experiment ruled out the possibility that the gradient in the tone sequence task arose from mental associations between movements and sounds, and revealed that MEG signal fluctuations unrelated to discrete sequential information can produce a pattern that resembles a gradient but cannot fully explain the gradient during the preparation of sequences of movements or tones. These results establish the CQ primacy gradient as a neural signature of serial order coding exhibiting at least partial domain-generality.

### 4.1 The CQ primacy gradient in perception

Our main finding reveals a clear CQ gradient in the auditory domain when participants prepared to listen to a familiar sequence of tones. This finding provides the first, to our knowledge, direct neurophysiological evidence that the CQ principle, previously established in the motor domain [13,14], might also govern serial organization in sensory domains, specifically auditory sequence memory. This parallel representation mirrors the CQ gradient we and other researchers observed in the motor domain [13,14].

Our hypothesis that a CQ principle underlies serial processing across several domains was motivated by behavioral evidence [3,4] and theoretical modelling across several domains [7,8,10,11], including the perception of melodies [9]. While direct neurophysiological evidence for CQ in non-motor domains has been lacking, other neural signatures for encoding of abstract serial order of auditory and visual sequences have been reported before. For example, the rank-order selectivity of single neurons, which has been identified in monkeys preparing for upcoming reaches [24,25] has also been identified for auditory, visual [26,27], and abstract sequences [28]. Similar rank-order selectivity was recently reported in hippocampal and entorhinal neurons of humans during spatial navigation [29] and presentation of visual sequences [30]. Together with evidence for separate encoding of sequence identity and serial position in human working memory [31–33], our finding of a neural signature compatible with a primacy gradient points to a plausible mechanism, namely Competitive Queuing, through which serial order emerges in behavior across motor and perceptual tasks, possibly reflecting a common neural architecture for sequencing information across domains.

In Experiment 2, we addressed a potential concern regarding the tone sequence task in Experiment 1, namely that the parallel gradient observed during the tone sequence task might reflect motor preparation, even in the absence of overt movement. Participants practiced the motor sequences in association with tones, each uniquely associated with a keypress. Numerous studies with a variety of methods have found engagement of motor areas when participants are passively presented with sensory effects they have previously associated with motor actions but not to stimuli that had not been associated with movement [22,23,34,35]. Thus, in our paradigm in Experiment 1, anticipating a tone sequence that has been associated with a motor sequence may automatically evoke motor-related activity, even when the task does not require any motor action. To exclude this possibility, in Experiment 2 participants listened to tones that had not been associated with any action. Experiment 2 thus provides evidence of a primacy gradient in a purely perceptual task.

### 4.2 The CQ primacy gradient in the motor domain

Our finding of a gradient during the preparation of a motor sequence replicates the main finding by Kornysheva et al. [14]. However, we did not reproduce the distinction between within-and across-sequence classifiers observed in their study. Kornysheva et al. [14] found a stronger gradient when decoding within-sequences, compared to across-sequences, pointing to the presence of effector-specific information both during execution and preparation of a motor sequence. In our study, decoding accuracies during sequence execution decreased from within-to across-sequence decoding, but Gradient Strength during sequence preparation did not (**Fig 4G bottom**, compared to **Fig 4A bottom**). While effector-specific information was thus clearly present during sequence execution in both studies, several differences in the acquisition method, and task design, between our study and Kornysheva et al. [14] could explain why we found no evidence of effector-specific information during sequence preparation. For example, we used magnetometers, while Kornysheva et al. [14] recorded from gradiometers and used a more challenging task than in our study, due to the recombinations of finger orders with different rhythms, and the use of irregular rhythms (non-beat-based, non-metrical). Furthermore, the amount of training might also affect the presence of effector-specific information during preparation for a movement. Participants in Kornysheva’s et al. [14] study trained for two days before the MEG session, while we administered only a short (20-30 minutes) training session.

We observed that the CQ gradient extended into the inter-trial interval. While Kornysheva et al. [14] did not report any quantification of the Gradient Strength during baseline, visual comparison of the figures between the studies reveals a more pronounced baseline gradient in our study (Figure 5A and Figure S4 in Kornysheva et al. vs. **Fig 4A,G bottom**, here). This difference might be attributable to variations in the predictability of visual cues across trials in both studies. In our design, the sequence of visual cues followed a more predictable pattern (two cues, alternating every three trials) compared to that used by Kornysheva et al. [14]. Combined with the lower difficulty of our task we described above, this higher predictability may have enabled participants to prepare for the next sequence already before the trial starts, although participants were not explicitly instructed to employ this strategy. We therefore calculated the baseline-corrected Gradient Strength as a more indicative measure of a preparation CQ gradient in the context of our paradigm. While some subjects might have predicted, either implicitly or explicitly, the next sequence during the baseline, the Gradient Strength significantly increased from the baseline to the preparation period.

### 4.3 Disentangling serial order from sequence-unrelated signals

Given that sequences necessarily unfold over time, Experiment 2 addressed the possibility that the classifier might pick up systematic MEG signal fluctuations over the course of a trial that are unrelated to the sequences of keypresses or tones. Potential sources for such systematic signal fluctuations may include the offset of the visual cue [36], participants’ mental tracking of time, anticipation of feedback at the trial end, or preparation to respond, or not, in order to register deviant trials at the end of the tone sequence task. To exclude the possibility that the gradient observed in the tone sequence task reflected such trial-structure–related activity rather than sequence-related representations, we devised a control task that preserved the trial structure but removed any perceptual or motor sequence. A gradient in this control task similar to the one in the tone sequence task would have indicated that there is no additional, sequence-related information that explains the gradient observed in the tone sequence task.

We observed above-chance classification of the first, second, and fifth “positions” in this control task, along with a pattern resembling a gradient, although this pattern displays four marked differences from the gradient observed in the tone sequence task and motor sequence task. First, the control task’s gradient was significantly less prominent than in the tone sequence task in Experiment 2. Second, Gradient Strength in the control task did not increase from baseline, unlike Gradient Strength in all sequential tasks in our study. This is in line with the idea that a full-fledged CQ gradient appears once participants have all the information to prepare for the specific upcoming sequence, consistent with previous findings [14]. Third, the change from baseline correlated between the two sequential tasks in Experiment 1, but it did not correlate between the tone sequence task and control task in Experiment 2. Fourth, the spatial topography of decoding accuracies, and of baseline-corrected Gradient Strength, differed significantly between the visual control task and the sequential tasks, hinting at different neural generators of each pattern. In sum, the CQ gradient observed in the sequential tasks in our study cannot be fully explained by trial-structure-related activity.

The neural pattern that resembles a CQ gradient in the control task may co-exist with the serial order code we observe during preparation for auditory or motor sequences. We therefore emphasize the importance of including appropriate control conditions when investigating CQ-related decoding results. Other researchers have addressed similar concerns when investigating the rank-order selectivity of single neurons in the macaque frontal cortex [37]. Their control tasks indicated that confounds such as tracking the passage of time and reward anticipation contributed, but did not fully explain, the observed pattern of results, which were therefore interpreted as an indication of true rank-order selectivity of neurons.

### 4.4 Domain-generality of CQ

Our goal was to test for neural evidence of the domain-generality of CQ. Reviewing the domain-generality of working memory, Nozari & Martin [38] argued that domain-generality can refer to a computational level, a neurophysiological or anatomical level, and a training or transfer level. In their view, evidence for the existence of domain-generality at one level does not necessarily entail domain-generality at another level. For example, computational equivalence does not imply neural equivalence, nor does neural equivalence imply computational equivalence. The serial order memory literature points to domain-generality of CQ at a computational level, as proposed by Hurlstone et al. [3]. Indeed, behavioral phenomena like the primacy and recency effects, or transposition errors, point to common principles of serial order coding across domains. However, other researchers argue in favor of domain-specificity in serial order coding, based on observations that the behavioral patterns of serial order working memory capacity across different domains can be better described by a model with independent, domain-specific serial order modules, compared to a model with a central, domain-general serial order module [39].

However, their argument in favor of domain-specificity does not necessarily conflict with our idea of domain-generality, as we use the term to highlight different aspects of serial order processing. Our usage of the term refers to a key computational aspect of CQ, the anticipatory, parallel, graded representation of sequence items. Even in the case of distinct serial order modules for each domain [39], this does not exclude the possibility that this key computational principle is shared by otherwise independent computations within each domain. Thus, the term domain-generality, in such a case, would point to structural similarities of architectures of the different domains. In this regard, our study aligns with Hurlstone et al. [3] in identifying whether shared computational mechanisms operate across different domains. To what extent such computational-level similarity across domains coincides with overlapping neural circuits for serial order coding, or translates into transfer potential of training, remains to be seen.

Our searchlight analysis provides evidence for topographic differences across domains at the sensor level, with right lateral frontal and temporal sensors showing a more pronounced baseline-corrected Gradient Strength for the tone sequence task than for the motor sequence task (**Fig 6**). At the same time, baseline-corrected Gradient Strength correlated across tasks in Experiment 1, suggesting that there are individual differences that drive the strength of the primacy gradient in more than one domain. Taken together, topographic differences, and inter-task correlation, are compatible with a partly domain-general, partly domain-specific mechanism of serial order coding at the neurophysiological level, while the underlying computational operations show structural similarity (the existence of a primacy gradient in both domains). This view is strongly aligned with the observations of Nozari & Martin [38] on working memory domain-generality.

### 4.5 Limitations and future directions

We extend the CQ gradient previously reported in the motor domain to the auditory domain. Further studies are needed to test the domain-generality of this gradient across other non-motor domains (visual, spatial, verbal etc.).

An important limitation in our study concerns the analysis of behavioral errors. The computational models for serial order processing are evaluated by how well they can simulate behavioral performance, including the types of errors encountered in serial order tasks [3,4]. Therefore, a neural signature of computational mechanisms should also be able to explain errors, ideally at the trial level. Our study does not lend itself to such exploration, due to the low difficulty level of our tasks and, consequently, the low error rates, leaving only a handful of trials eligible for analyzing behavioral error patterns. Different paradigms with more difficult tasks and/or different behavioral readouts of serial order processing (as in, for example, [16]) may elucidate the relationship between serial order errors and neural CQ gradient. Lastly, single-trial neural signatures are paramount for explaining errors. Kornysheva et al. [14] explored the relationship between neural gradient and behavior, and found a significant correlation at the inter-individual level, as well as on a trial-by-trial basis within participants, but only for 5 out of 16 individuals. The non-ubiquitous within-subject correlation with behavior could have partly resulted from the exclusion of all the trials with incorrect keypresses in their analysis, and thus the quantification of the neural gradient derived only from the remaining, correct trials. In any case, it is reassuring that at the inter-individual level and at the trial-wise within-individual level for some individuals, a negative correlation between behavioral error and Gradient Strength was found (note that the authors name their dependent variable probability distance, which is methodologically identical to Gradient Strength in our study). Ideally, future studies should focus on study designs that allow a link between behavioral performance and neural signature at the single-trial level.

Another limitation stems from our methods of analyzing the neural data. Our methods (see also [13,14]) rely on averaging across trials and/or time bins in order to reach the final metric of Gradient Strength. This renders the method blind to any transient activity that is not time-locked to cue onset/offset, such as sequential rather than parallel pre-activations, e.g. replays during ripples. As a consequence, as already pointed out [40], we cannot determine whether incomplete sweeps of neural replay during retrieval of the cue-associated sequence may give rise to the CQ gradient we observe after averaging, or whether there is any replay that independently coexists with the CQ gradient. While such independent coexistence is not unlikely in a recent synaptic model of serial order encoding in working memory [41], future studies should elucidate the relationship between replay and the CQ gradient by using mixed methods in the same experiment (decoding of replay, e.g. as in [42], and signatures of graded representations, e.g. as in our study or the neural manifold geometries of item-order multiplexing in [43]).

### 4.6 Conclusion

In summary, we extend an essential principle of serial order coding in the brain, the parallel anticipation of upcoming serial events, from movement planning to perception. Our study thus provides evidence for a partial domain-generality of the neural mechanisms underlying sequential ordering, and lends support to the primacy gradient as a key computational principle of a mechanistic serial order model, Competitive Queuing. We put our neurophysiological evidence of this gradient to a rigorous test, and rule out potential contributions of systematic, sequence-unrelated neural activity. Our study thus reveals a de-confounded and domain-general signature of serial order coding in the human brain within the framework of Competitive Queuing.

## 5 Methods

### 5.1 Participants

We recruited 50 healthy adult participants (22 females, aged 20 - 44, mean age = 27.9 ± 5). Participants were right-handed, assessed with the Edinburgh Handedness Inventory, had normal or corrected-to-normal vision, and reported no history of psychiatric or neurological conditions or impairments of hand function. None of the participants was a professional musician. Eleven participants had an amateur musical background (mean years of experience = 4.6 ± 3.8, range: between 1 and 10). The other thirty-nine participants reported no musical background. We recruited participants via Sona Systems (https://www.sona-systems.com) and the MEG lab’s contact list of participants from previous experiments. The study was approved by the ethics committee at the Medical Faculty at Otto-von-Guericke University Magdeburg. Participants provided written informed consent and received financial compensation for their time.

We excluded two participants due to poor performance in the motor task during the MEG session (41% and 36% Error Rate, respectively, i.e., percentage of trials with incorrect finger presses), one participant due to a technical error of the response device, and one participant due to excessive artifacts in MEG. We report data from 46 participants, half of whom took part in Experiment 1 (9 females, aged 20-44, mean age = 26.6 ± 5.3), and half in Experiment 2 (i.e., none of the participants of Experiment 1 took part in Experiment 2; 11 females, aged 22-42, mean age = 29.1 ± 4.4).

### 5.2 Apparatus

During MEG recording, participants used a left-hand response device (LUMItouch Response System; Photon Control Inc.) with five keys. Participants operated the device with their left (non-dominant) hand for producing sequences of keypresses (all five fingers of the left hand), and with their right hand for reporting deviants in the tone sequence task (only right-index finger). Auditory stimuli were presented via MEG-compatible plastic tube earphones. All auditory stimuli were loudness-matched at -18 Loudness Units Relative to Full Scale. We adjusted the sound pressure level for each participant to a subjectively comfortable level before the experiment. A projector presented visual stimuli on a screen, approximately 100 cm in front of the participant. The screen background was black.

For practice sessions before the MEG session, participants used desktop computers outside the MEG chamber (Lenovo ThinkCentre M70s, with 24” ASUS VG248 monitor), equipped with standard PC keyboards (Logitech K120). During practice, we provided auditory stimulation via headphones (AKG K371), with the volume individually set at a comfortable level.

The experiment was scripted with Psychtoolbox version 3 [44] running in MATLAB version 9.12.0 (R2022a) [45].

### 5.3 Task and procedure

#### 5.3.1 Experiment 1

Experiment 1 aimed to reveal whether preparation for an upcoming sequence of tones entails a similar, parallel representation of serial order as previously observed for sequences of movements. To this aim, participants were tested on two tasks, a motor sequence task, and a tone sequence task (**Fig 1A, B**).

Before the MEG session (see below), participants learned two sequences of five finger movements, each associated with a distinct fractal cue; every keypress produced a tone according to a fixed key–tone mapping (notes C–G). Thus, each participant learned two motor sequences and two corresponding tone sequences. From a set of four fractals, two fractals were randomly selected for each participant and each fractal associated with one specific sequence of finger presses and its corresponding sequence of tones. The order of keypresses in each of the two sequences was determined randomly for each participant, with the following constraints. In each sequence, each of the five keys of the response device had to be pressed once. Neither sequence could include ascending or descending triplets of keypresses of neighboring keys, such as index finger – middle finger – ring finger, or the reverse. In addition, the two sequences had to start with the same key (adapted from Kornysheva et al. [14]), but could not share any other keypress in the same position. Finally, the two sequences could not share any transitions between consecutive keys. In addition, each keypress triggered one tone according to a fixed key-to-tone mapping. Specifically, from left to right (little finger to thumb), the five keys produced the five tones C, D, E, F, and G, respectively (sine-wave tones at frequencies 261.63, 293.66, 329.63, 349.23, and 392 Hz). Each tone lasted 200 ms, including a 10 ms ramp at tone onset and offset.

During MEG recording, participants completed two tasks in separate blocks of trials. In the motor sequence task, participants produced the two learned sequences from memory under two conditions: either performing the keypresses silently (“*silent production”* condition; **Fig 1A**), where keypresses did not trigger any tones, or producing the associated tones (“*production with tones”* condition; **Fig 1 in S1 Supporting Information**). In the tone sequence task, they listened to the corresponding sequences of five tones and identified deviations from the memorized patterns (**Fig 1B**).

In both tasks, the fractal at trial onset informed participants which specific sequence of movements, or tones, to prepare (**Fig 1**). Fractals were presented at the center of the screen, inside a white contour of a left hand (visual angle approximately 20°) with white discs covering the fingertips. Between 1.8 and 2.2 s after cue onset (uniform random distribution), the cue disappeared, while the contour of the hand remained on the screen. Analyses focused on this preparatory period, before the onset of the first movement or tone, to examine representations of serial order during sequence planning. The offset of the cue signaled the beginning of the response window for the motor sequence task (Go cue), and the beginning of the listening period for the tone sequence task, each lasting 3.1 s. From cue offset onwards, events differed between the two tasks and are therefore described separately.

### Motor sequence task

After fractal offset, participants had to produce the correct sequence of five keypresses at a pace of 2 Hz, starting 500 ms after cue offset (**Fig 1A**). Following the response window, participants received visual performance feedback regarding accuracy of keypress order, and precision of keypress timing. Specifically, five white horizontal lines represented the five presses, and a gray fixed dashed line along the horizontal meridian of the screen represented zero deviation from the target timing (**Fig 1A**). The distance of each keypress line from the dashed line was scaled to represent the timing error of the corresponding keypress relative to its target timing (placed above for a late press and below for an early press). For motivational purposes, participants earned points (ranging from 0 to 4) for precise timing, measured as the mean absolute press time deviation in each trial, and saw a running sum of points earned so far at the end of each trial. An incorrect keypress turned the respective line to red color and nullified any points for that trial. The performance feedback screen lasted 2 seconds, followed by a jittered inter-trial interval (ITI) between 0.8 and 1.2 s (black screen).

In trials in which participants pressed any key during preparation, i.e., while the visual cue was still on the screen, they received a warning that they pressed too early and should wait for the Go cue (i.e., the disappearance of the fractal), and the trial was restarted. In trials in which they produced less than, or more than five keypresses, participants received a warning that they pressed too few, or too many keys, respectively. These trials were excluded from analysis.

### Tone sequence task

The tone sequence task (**Fig 1B**) closely matched the motor sequence task with respect to event timing and sequence design. However, participants had to refrain from any movement during the listening window. Instead, they heard one of the two sequences of five tones, whose order was predicted by the visual cue at trial onset. Thus, tones were played in the absence of any keypress, at a pace of 2 Hz, with the first tone played 500 ms after cue offset. Any keypress during the listening window triggered a warning that participants should not move during the listening window of the tone sequence task, and the trial was aborted and restarted.

To ensure that participants attended to the visual cue at trial onset, and prepared for the corresponding tone sequence, they had to detect rare deviant trials. There were two types of deviant trials. In *sequence cue mismatch trials* (8.33% of all trials in the tone sequence task), the mapping between visual cue and tone sequence was unexpectedly reversed (for example, the visual cue for sequence A was presented but followed by tone sequence B). This type of deviant trials thus enforced attention to the mapping between the visual cue and the subsequent tone sequence. In addition, we ensured that participants attended to the tone sequences until the last tone was played, by including *last tone mismatch trials* (8.33% of all trials in the tone sequence task, not coinciding with *cue mismatch trials*). In these trials, the mapping between visual cue and tone sequence was as expected, however, the last tone in the sequence was a repetition of one of the first four tones. To minimize guessing and false alarms, we took care of making the last tone deviance as salient as possible by maximizing the difference from the last tone transition. On each trial, the listening window was followed by the word “Mismatch?”, presented in white font on the screen until a response was made or for a maximum of 2 seconds. If participants wanted to report that the current trial was a deviant trial (of either deviant type), they had to press a key with their right hand index finger within 2 seconds, or else refrain from pressing that key. We excluded all trials with such a response, as well as all remaining, undetected deviant trials, from all analysis.

Trials ended with visual and auditory performance feedback. Participants heard a success sound for all hits and correct rejections in the deviant detection task, and an error sound for all misses and false alarms. In addition, they earned 2 points per correct decision and lost 20 points for each incorrect decision, and saw on the screen a running total of points earned in the tone sequence task so far. This visual performance feedback lasted 1 second, and was followed by an inter-trial interval between 0.8 and 1.2 s (black screen).

### Design

During MEG recording, participants completed 18 blocks, organized in cycles of three blocks each. Each cycle consisted of one block of the tone sequence task and two blocks of the motor sequence task, one for each of the “*silent production*” and “*production with tones*” conditions. We randomized the order of blocks within each cycle, and informed participants about the upcoming task before each block.

Each block consisted of 24 trials. The same sequence was cued for three consecutive trials before a switch to the other sequence. This yielded a total of 144 trials for the tone sequence task, and for each condition of the motor sequence task. In the tone sequence task, each block contained four deviant trials, two of each type, randomly placed within the block. Blocks were separated by short breaks (around 40 seconds). The MEG recording lasted approximately 75-80 minutes.

### Training session outside the MEG

Before the MEG recording, participants completed a training session outside the MEG chamber, which lasted approximately 30-40 minutes. The aim of this session was for participants to learn to produce the two sequences of keypress from memory, as instructed by the visual cues at trial onset, and to learn the corresponding two tone sequences generated by the fixed key–tone mapping. The training session consisted of two steps.

First, participants completed at least two blocks of a keypress synchronization task, whose structure was virtually identical to the motor sequence task (*production with tones* condition), with the exception that we visually cued each individual keypress during the response window. Cues for individual keypresses were green discs, presented at the fingertip of the displayed hand contour. Each green disc instructed a keypress with the corresponding finger at the exact time at which the green disc appeared. Correct keypresses within ± 60 ms of the onset of the green disc turned the disc’s color from green to purple. Purple discs granted bonus points (1 per disc) on top of the timing precision points, displayed separately during the feedback screen. After the two blocks, we asked participants whether they felt confident in producing each of the two sequences from memory. If they indicated that they required more training, we repeated the keypress synchronization task until they felt confident to move on to the second training.

In the second training, one third of trials were identical to the keypress synchronization task described above (every third trial, starting from the first trial). In the remaining trials, there were no visual cues during the response window, i.e., participants completed the sequence from memory, based on the fractal presented at trial onset. However, to maintain participants’ motivation, each correct and temporally precise keypress still triggered the presentation of a purple disc on the corresponding fingertip on the screen, as described above. Importantly, during MEG recording of the main experiment, we provided no visual stimulation (neither visual cues nor visual feedback in the form of purple discs) during the response window of the motor sequence task, nor during the listening window of the tone sequence task.

Participants whose performance was poor (i.e., a large number of trials with incorrect finger presses) after two blocks of the second training step completed additional blocks until they felt confident. On average, participants completed 4.72 ± 1.09 training blocks in total (range: between 4 and 8). Training took place either immediately before the MEG session (12 participants), or on the previous day (11 participants).

Since the objective of Experiment 1 was to replicate previous findings with a motor sequence task [14] and to extend these findings to the auditory domain when participants anticipate an associated sequence of tones, we did not have any *a priori* hypothesis regarding the comparison of the two motor task conditions *(silent production* vs. *production with tones*). Thus, for brevity, we focus on the *silent production* condition. Results from *production with tones* condition were found to be similar to the *silent production* and are reported in the Supporting Information (**Fig 1 in S1 Supporting Information**). In the results and discussion, every mention of the motor task refers to the silent production condition, unless otherwise noted.

#### 5.3.2 Experiment 2

The goal of Experiment 2 was to replicate our finding in Experiment 1 that preparation for an upcoming sequence of tones entails a similar, parallel representation of serial order similar to that observed for upcoming movement sequences. In addition, Experiment 2 aimed to rule out two key alternative interpretations.

First, given that each fractal in Experiment 1 was associated with a sequence of tones, as well as a sequence of movements, Experiment 1 could not rule out the possibility that visual cues triggered (implicit) preparation of a movement sequence even in the tone sequence task, without any eventual, overt movement during the listening window. To rule out this possibility, we dropped the motor sequence task in Experiment 2, and included only the tone sequence task, which was identical to Experiment 1 (**Fig 1B**). A new cohort of naïve participants was recruited, ensuring that they never associated the visual cues with any motor sequence.

Second, in Experiment 1 an alternative explanation may account for the decoding of order information from MEG data. Given that sequences unfold over time, a classifier meant to discriminate between sequence positions may in fact reflect time-dependent MEG signal fluctuations that are unrelated to serial order. For example, we expected the MEG signal to vary systematically over time following the offset of the visual cue. To control for a bias in classification due to such sequence-unrelated information, Experiment 2 included a control task whose trial structure was identical to the tone sequence task until the end of the listening window, except that no tones were presented following the visual cue (**Fig 1C**). To ensure that the visual cue could not trigger any representation of any previously experienced sequence, participants first completed the control task before learning the mapping between visual cues and tone sequences for the subsequent tone sequence task. To minimize differences in task engagement between the control task and the tone sequence task, the control task included a delayed match-to-sample task. At the onset of each trial, we presented one of the two fractals used as visual cues in the tone sequence task (same duration, alternating every three trials). Following fractal onset, there was a delay window of 3.1 s, followed by a second fractal, which could be identical to the first (in 83.33% of trials) or not. In the latter case, the second fractal was randomly chosen from the set of four fractals used as visual cues in the tone sequence task across participants. The task was to memorize the first fractal and press a key with the right-hand index finger if the second fractal was different from the first. Similar to the tone sequence task, participants had 2 seconds to respond, followed by similar feedback as in the tone sequence task. In sum, the control task involved the same visual stimulation and overall trial structure, and required similar task engagement, as the tone sequence task, but abolished any serial order information.

Participants first completed six blocks of the control task, yielding 144 trials in total. Following the control task, participants completed two blocks of the tone sequence task (without the deviation detection task and without any deviant trials) in order to learn the mapping between the two visual cues and the two tone sequences. Participants were instructed to learn the sequence of tones and their association with the visual cues through observation. After this training, every participant felt confident remembering the mapping of visual cues and tone sequences, and moved on to six blocks of the tone sequence task (144 trials), which was identical to the tone sequence task in Experiment 1. Due to the short training period (2 blocks), we reduced the penalty of an incorrect decision in the mismatch task from 20 points to 2 points. The same penalty applied for the visual control task. We recorded MEG during the control task and the tone sequence task, which, together, took approximately 60 minutes.

### 5.4 MEG recording

We recorded MEG data continuously via the Elekta Neuromag TRIUX triple sensor system with 102 magnetometers and 204 gradiometers (Elekta Neuromag, Oy, Helsinki) inside a µ-metal-shielded room (Vacuumschmelze, Hanau, Germany). We asked participants to minimize movement during MEG recording, especially head movements and blinks. Small foam cushions helped participants to hold their heads in place.

Simultaneous to the MEG, the electrooculogram (EOG) was recorded via three electrodes placed at the outer canthi of both eyes (for horizontal eye movements) and beneath the right eye (for vertical eye movements). In addition, we recorded surface electromyography (EMG) via electrodes placed on the skin of the left hand to capture activity of the abductor polices brevis muscle, the abductor digiti minimi muscle, and the first dorsal interosseous muscle with a belly-tendon montage, and the flexor carpi radialis muscle with a belly-belly montage.

We sampled MEG, EMG, and EOG data at 1000 Hz. MEG data were bandpass-filtered online between DC and 330 Hz, and EMG/EOG data bandpass-filtered online between 0.1 and 330 Hz. Prior to the recording, we digitized participants’ three-dimensional skull shape and individual landmarks (nasion, left and right preauricular point) in the 3D space with a Polhemus digitizer (Polhemus 3SPACE FASTRAK system; Polhemus, Colchester, VT, USA). During the experiment, we recorded head position intermittently during the breaks (between trial blocks, i.e. every 24 trials), via five Head Position Indicator (HPI) coils (inion, nasion, left and right preauricular point, and vertex) placed on an EEG cap (Easycap, Herrsching, Germany).

### 5.5 Behavioral data analysis

For the motor sequence task, we defined three performance measures. We examined whether participants expanded or compressed temporal intervals between consecutive keypresses by calculating the Inter-Press Interval (IPI; median across intervals and trials). We quantified Timing Variability as the interquartile range of the IPIs (computed across intervals and trials). For IPI and Timing Variability only trials with correct finger presses were included. Finally, we defined an Error Rate as the percentage of trials that contained erroneous key presses (wrong finger). Invalid trials (more or less than 5 presses) also counted as errors. Performance in the tone sequence task and the control task were evaluated using the sensitivity index (d’) [46]. Hit rates and false alarm rates at exactly 1 or 0 were adjusted to 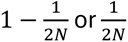 respectively, where N is the total number of trials for each task, i.e. 144 (6 blocks x 24 trials).

### 5.6 MEG data analysis

We analyzed MEG and EMG data in Matlab [45], using the FieldTrip toolbox (version 20240515; http://fieldtriptoolbox.org) [47] and custom MATLAB code. The script for Linear Discriminant Analysis (LDA) was the script used in Kornysheva et al. [14], found in https://data.mendeley.com/datasets/fxbmm66cr6/1.

#### 5.6.1 Pre-processing

Data were Maxwell filtered (MaxFilter Software, Elekta Instrument Ab Stockholm, Sweden). Maxwell filtering included signal space separation (SSS) to suppress spatial artifacts originating outside the helmet, movement correction using the intermittent HPI recordings, and downsampling from 1000 Hz to 500 Hz. We report data recorded via the 102 magnetometers.

Pre-processing included attenuation of eye movement and heart artifacts via Independent Component Analysis (ICA). For the ICA, we created a high-pass filtered copy of the continuous data (windowed sinc finite impulse response (FIR) filter in FieldTrip: cutoff frequency 1 Hz, Hamming window, filter order 826, zero-phase), and cut the continuous data into epochs, starting from the beginning of the inter-trial interval to the beginning of the next inter-trial interval. We then performed ICA in FieldTrip (runica), and identified components that corresponded to blinks, other eye movements, and heart artifacts, based on the topography of components, their power spectrum, and their coherence with the EOG. Obtained ICA weights were then applied to the original, continuous data, rejecting the components that captured artifacts. Thereby, we avoided any spectral filtering of the data that we eventually analyzed.

Because the duration of the visual cue was jittered between 1.8 and 2.2 s, we created two separately epoched datasets from the ICA-corrected, unfiltered continuous data. One dataset was time-locked to the fractal offset [-2.6 s to 3.4 s] to align analyses to the Go cue, and the other to the fractal onset [-0.8 s to 5.2 s] to capture cue-related activity. Both datasets were baseline-corrected using the same pre-cue window, i.e. [-0.8 s to 0 s] relative to fractal onset, which corresponds to the longest possible ITI window that was available in all trials.

We removed all trials in the motor sequence task with less than, or more than, five keypresses, as well as trials with any erroneous keypress. In the tone sequence task as well as the control task of Experiment 2, we removed all deviant trials and all remaining trials with a response. Due to excessive head-movement-related artifacts, we excluded from the analysis one block of the motor sequence task in one participant, as well as one block and one trial of the motor sequence task, and four trials from the tone sequence task in another participant. These trials with excessive artifacts were excluded before the ICA procedure. All other excluded trials (erroneous keypresses, attentional task deviants, and false alarms) were initially included in the ICA but excluded from further analysis.

#### 5.6.2 Classification analysis

We aimed at investigating whether humans prepare several items of an upcoming sequence of tones in advance, relying on a parallel representation of serial order during preparation similar to that observed for movement sequences [13,14]. To address this question, we conducted a multivariate pattern analysis (MVPA) of the pre-processed MEG data, using LDA (adapted from Kornysheva et al. [14]). For each participant, we trained classifiers to discriminate between the five sequence items during production or listening, for each task separately. Training involved the mean MEG signal per sensor across a 100 ms time window before each keypress in the motor sequence task, and before each tone in the tone sequence task. For the control task in Experiment 2, which had no events during the delay window, we trained the classifier to discriminate between five time points during the delay window, defined as the 100 ms intervals preceding the five latencies at which tones were presented in the tone sequence task. For all classifiers, the features were the 102 magnetometer channels and the assigned class labels were each item’s ordinal position in the sequence. We set the regularization parameter to 0.01.

In the first step, we evaluated the LDA’s accuracy to distinguish the sequence items. A ‘within-sequence’ classifier was trained on items from trials belonging to one sequence and classified items from trials belonging to the same sequence, with a k-fold cross-validation (5 folds by shuffling trials, and 5 repetitions). Since the class labels reflect the ordinal position and each item appears exactly once in the sequence, there is a one-to-one relationship of ordinal position and effector/tone pitch identity. Therefore, the ‘within’ classifier picks up and decodes both positional and effector/tone pitch information. To isolate the positional information, an ‘across-sequence’ classifier was trained on items from trials belonging to one sequence, but classified items from trials belonging to the other sequence. Since this is a form of cross-decoding between different datasets, no cross-validation was necessary. There were two sequence runs for each classifier (‘within’: sequences A => A, and B => B; ‘across’: A => B, and B => A), resulting in one confusion matrix per sequence run. Confusion matrices were averaged across sequence runs for each classifier. We performed this classification procedure for each task separately.

In the second step (**Fig 3**), we tested for representations of the upcoming sequence during preparation, i.e., during the time window of the visual cue, following a similar approach to Kornysheva et al. (2019). We employed the ‘within’ and ‘across’ classifiers described above, using as test data the MEG activity at each sensor averaged over consecutive 10 ms time bins across the entire trial epoch, and decoded each time bin. The ‘within’ decoder was cross-validated as described above (5 folds by shuffling trials, 5 repetitions). As output of the decoder, we obtained the classification score for each class, i.e., five values per time bin. Following Kornysheva et al. [14], we transformed classification scores so that they ranged from 0 to 100 with the sum of all transformed scores amounting to 100, by subtracting the lowest score from all scores, dividing all scores by their sum, and then multiplying the result by 100. This transformation was carried out separately per time bin and per trial. The resulting metric, termed “pattern probabilities” in Kornysheva et al. (2019), indicates how likely it is for each time bin to be classified as each of the five classes, with one value per class. This procedure was performed in two sequence runs for each decoder and for each task.

To test whether these probabilities are ordered according to the rank of movements in an (upcoming) motor sequence, or the rank of tones in an (upcoming) sequence of tones, we followed the approach described in Kornysheva et al. [14]. We first calculated the median difference between probabilities of successive ranks (i.e., classes; 1^st^ press minus 2^nd^ press, 2^nd^ minus 3^rd^, 3^rd^ minus 4^th^, and 4^th^ minus 5^th^; equivalent for tones), a metric we term here Gradient Strength. This resulted in one value for each time bin and each trial. Given that these values were typically non-normally distributed, we computed their median across trials for each time bin. For the cross-validated ‘within’ classification, we took the median across trials for each repetition and then the mean across repetitions.

We then computed the mean across time bins in two time windows of interest, and then for each time window of interest the mean across sequence runs, in order to obtain a single quantification of a Competitive Queuing gradient, i.e. Gradient Strength, for each task and time window. The first time window spanned the last second before the offset of the visual cue [14]. We compared the Competitive Queuing gradient in this time window with the gradient in a second time window, spanning the window of the inter-trial interval that was available for all trials (from -0.8 to 0, relative to fractal onset). For illustration purposes, we also show the time-course of pattern probabilities baseline-corrected using each probability’s mean across the baseline window, i.e., the last 800 ms before cue onset (e.g. in **Fig 4A, top**; time axis relative to cue onset).

#### 5.6.3 Searchlight analysis

To identify which channels contribute most to the observed Competitive Queuing gradient, we performed a searchlight analysis. For this, we ran 102 searchlight iterations, one for each channel. Each iteration followed the cross-decoding analysis described above in step two, but included as features only the current channel of interest and its two closest neighbors, identified via their Euclidean distance in 3D space to the channel of interest. Each iteration resulted in one value per trial, i.e. Gradient Strength, as described above, for the time bin at 500 ms before cue offset. We then computed the median across trials. For the ‘within’ cross-validated output, we computed the median across trials for each repetition, and then the mean across repetitions, per channel. Finally, we averaged the across sequence runs per channel, for each decoder separately.

To enable comparisons between the preparation period (here 500 ms before cue offset) and the ITI baseline, we did the same analysis for the time window [-0.8 to 0 s] relative to fractal onset, with the added step of averaging across time bins after taking the median across trials. This resulted in two values of Gradient Strength per channel and decoder, one for each time window of interest. Since we found a significant positive difference during the ITI, we subtracted the Gradient Strength of the ITI from the Gradient Strength of the preparation period per channel, which we display as a topography. This topography reveals the baseline-corrected Gradient Strength for each channel. We conducted the searchlight analysis separately for each task.

Similarly to the first step of the main analysis, we investigated which channels show the highest decoding accuracies when discriminating sequence items (102 iterations, one for each channel and its two closest neighbors). The ‘within’ classifier was trained on sequence items of one sequence and classified items of the same sequence (with a k-fold cross-validation; 5 folds by shuffling trials, 5 repetitions), and the ‘across’ classifier was trained on items of one sequence and classified items of the other sequence. We obtained as an output decoding accuracies. After averaging across sequence runs (and folds and repetitions for the ‘within’ classifier), this resulted in one value per channel, which we display as a topography. We performed this analysis separately for each task.

### 5.7 Statistics

To evaluate the training of the classifier, we used Monte-Carlo simulations at the subject-level. Specifically, we constructed a null distribution of decoding accuracies obtained by shuffling the class labels of the training dataset (n = 100000 permutations) and compared the actual decoding accuracy (from real labels) to the null distribution. The empirical means of the null distributions were found to be, at the group average, at 20% ± 0.008%. *P*-values were computed as the percentage of the shuffled accuracies being higher than the actual accuracy. Decoding accuracy was considered statistically significant when this *p*-value was below 0.05. 21 of 23 participants showed significant decoding accuracy in the motor sequence task in Experiment 1; all participants in Experiment 1 and all participants in Experiment 2 showed significant accuracy in the tone sequence task. A group-level analysis involved the confusion matrices from the first step of the main classification analysis, by testing whether each element of the diagonal is above the chance level (20%) at the group level with one-sided one-sample t-tests, Bonferroni-Holm corrected for five comparisons. This was done for each classifier and task separately.

Regarding the gradient quantification, i.e. Gradient Strength, we expected a more pronounced CQ gradient in the last second before cue offset, compared to the ITI, both in the motor sequence task and the tone sequence task, but not in the control task. To test this, we conducted a 2 x 2 x 2 repeated-measures ANOVA on Gradient Strength in Experiment 1, which included the three within-subject factors Decoder (‘within’, ‘across’), Phase (ITI, last second before cue offset) and Task (tone sequence task, motor sequence task). We expected a significant main effect of Phase, as well as Decoder, following Kornysheva et al. [14]. For Experiment 2, we computed a similar 2 x 2 x 2 repeated-measures ANOVA, which included the three within-subject factors Decoder, Phase and Task (tone sequence task, control task), and expected a significant Phase x Task and Decoder x Task interaction. Additionally, to investigate the presence of the hypothesized gradient, we tested for a positive Gradient Strength by performing one-sided one-sample t-tests against zero for each Decoder, Phase, and Task levels, Bonferroni-Holm corrected for a family of eight comparisons for each experiment. We also tested whether the Gradient Strength during the last second before cue offset was larger than the Gradient Strength during the ITI with planned comparisons (two-sided paired-samples t-tests) Bonferroni-Holm corrected for a family of four comparisons for each experiment separately.

To investigate whether participants who show a pronounced gradient in one task also do so in the other task, we tested for Pearson’s correlation across tasks using the baseline-corrected Gradient Strength, separately for each Experiment. A statistically significant correlation between tasks of Experiment 1 together with a non-significant correlation between the tone task and the control task for Experiment 2 would provide further evidence for a domain-general representation of serial order.

We ran all statistics on Gradient Strength in JASP version 0.19.3 [48].

Searchlight topographies of baseline-corrected Gradient Strength were tested for differences across tasks, with a paired-samples 2-tailed t-test, with a Monte-Carlo cluster-based multiple comparisons correction (cluster forming threshold *p* = 0.05, n = 10000 permutations, alpha = 0.025 per tail) [49], separately for each decoder and experiment. We ran the test using Fieldtrip.

## 6 Acknowledgments and funding sources

We would like to thank Maria Kaiser and Leonie Orlikowski for their contributions to data collection.

## 8 Supporting Information

S1 Supporting Information. Additional data from *production with tones* condition and control analysis to rule out EMG-related activity.

## Notes

https://doi.org/10.5281/zenodo.18457101

### Competing Interest Statement

The authors have declared no competing interest.

