## Supporting Information 1 for "A domain-general neural signature of serial order memory across action and perception"

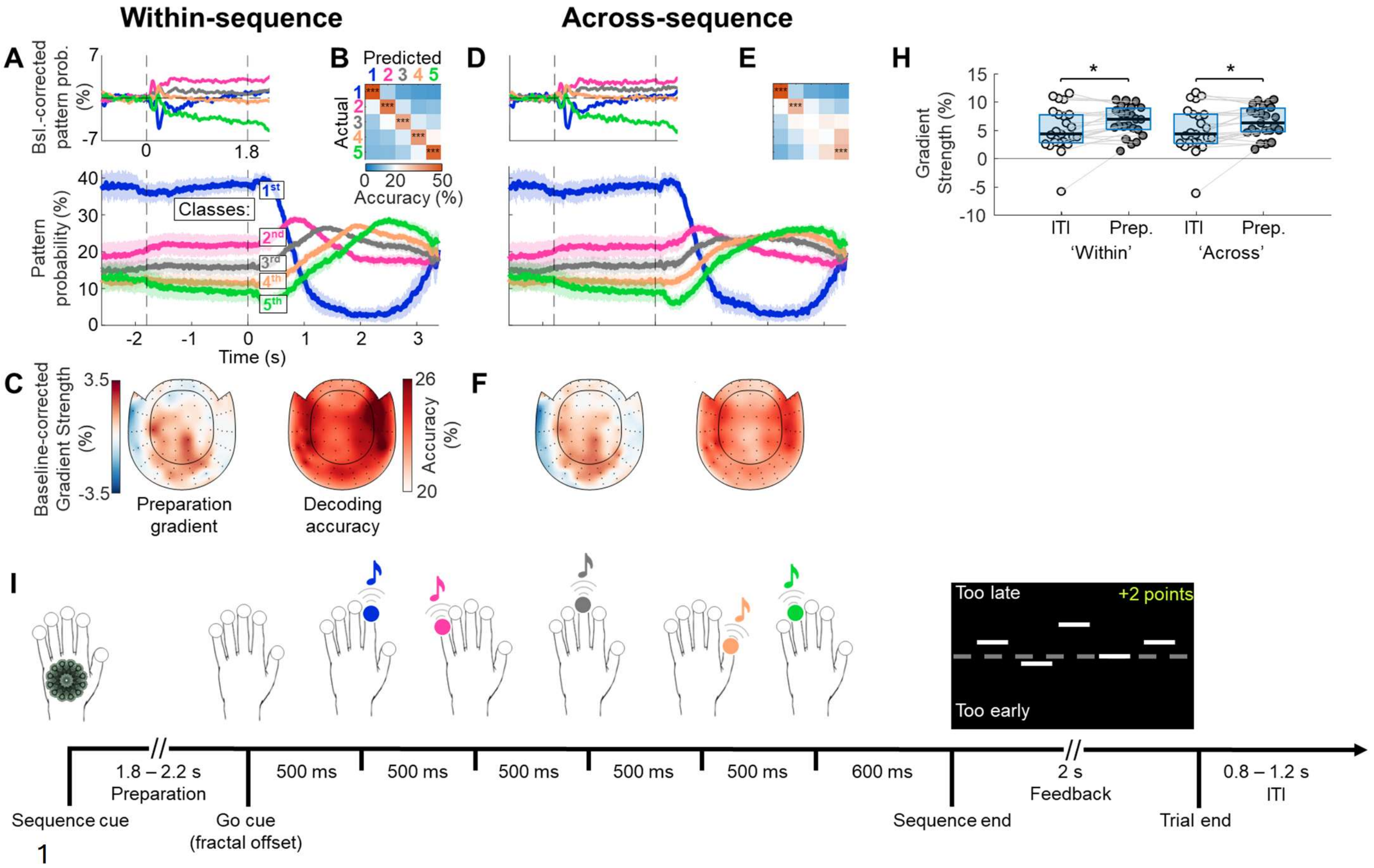


**Fig 1. Results from the *production with tones* condition are similar to the *silent production* condition.** During the MEG recording session, participants also performed a condition of the motor task where each key press triggered a tone **(I)**, identical to the condition they experienced during the behavioral practice session. We present here the results from this condition following the same analysis steps as in the analysis of the silent motor production condition. Qualitatively, the results are very similar to the silent motor production condition, including all the statistical inference. **A-C,** within-sequence decoding (sequence A=>A, and B=>B), 5-fold cross-validated. **A, lower panel** shows the grand-average time-course of pattern probabilities for each of the five classes, with color coding for class, i.e., position in the sequence. The time axis is time-locked to cue offset, which signaled the start of the presentation window in the tone sequence task. The dashed line at t = -1.8 s indicates the latest possible cue onset. Shadings represent 95% confidence intervals. **A, upper panel,** difference in pattern probabilities from baseline (-0.8 to 0 s relative to cue onset), time-locked to cue onset. The dashed line at t = 1.8 s indicates the earliest possible cue offset. In both panels of A, the time window enclosed by the dashed vertical lines is the shortest possible window of sequence preparation. Time-courses in panel A, upper, are smoothed with a Gaussian window spanning five bins (=50 ms). **B,** group average confusion matrix of the classifier's cross-validated decoding performance to discriminate items of the same sequence. Elements of the diagonal are tested against chance level (20%), and *p*-values are

Bonferroni-Holm corrected for five comparisons. **C,** searchlight results, indicating which sensors show the highest baseline-corrected Gradient Strength (left), and which sensors show the highest decoding accuracy when classifying sequence items during the presentation window (right). **D-F,** same as A-C, but for across-sequence decoding (sequence A=>B, and B=>A). **M,** Quantification of the degree to which pattern probabilities are graded by rank. The y-axis shows the Gradient Strength, i.e., the median difference of pattern probabilities between successive pairs of positions per participant, summarized across trials, sequences (and repetitions for 'within' decoding), and time bins from two time windows of interest: the baseline ('ITI') and the last second before cue offset ('Prep.'). Gradient Strength was significantly above zero even during the ITI, possibly due to the predictable switching of sequences in our paradigm which may have allowed participants to predict the upcoming sequence well before the cue onset. The box edges define the upper and lower quartiles. The horizontal black lines indicate the group median. We compared the Gradient Strength between the baseline and the last second before cue offset, for both tasks and decoders, with paired-samples t-tests ($p$-values Bonferroni-Holm corrected for four comparisons). * $p < 0.05$, ** $p < 0.01$, *** $p < 0.001$, Bonferroni-Holm corrected.

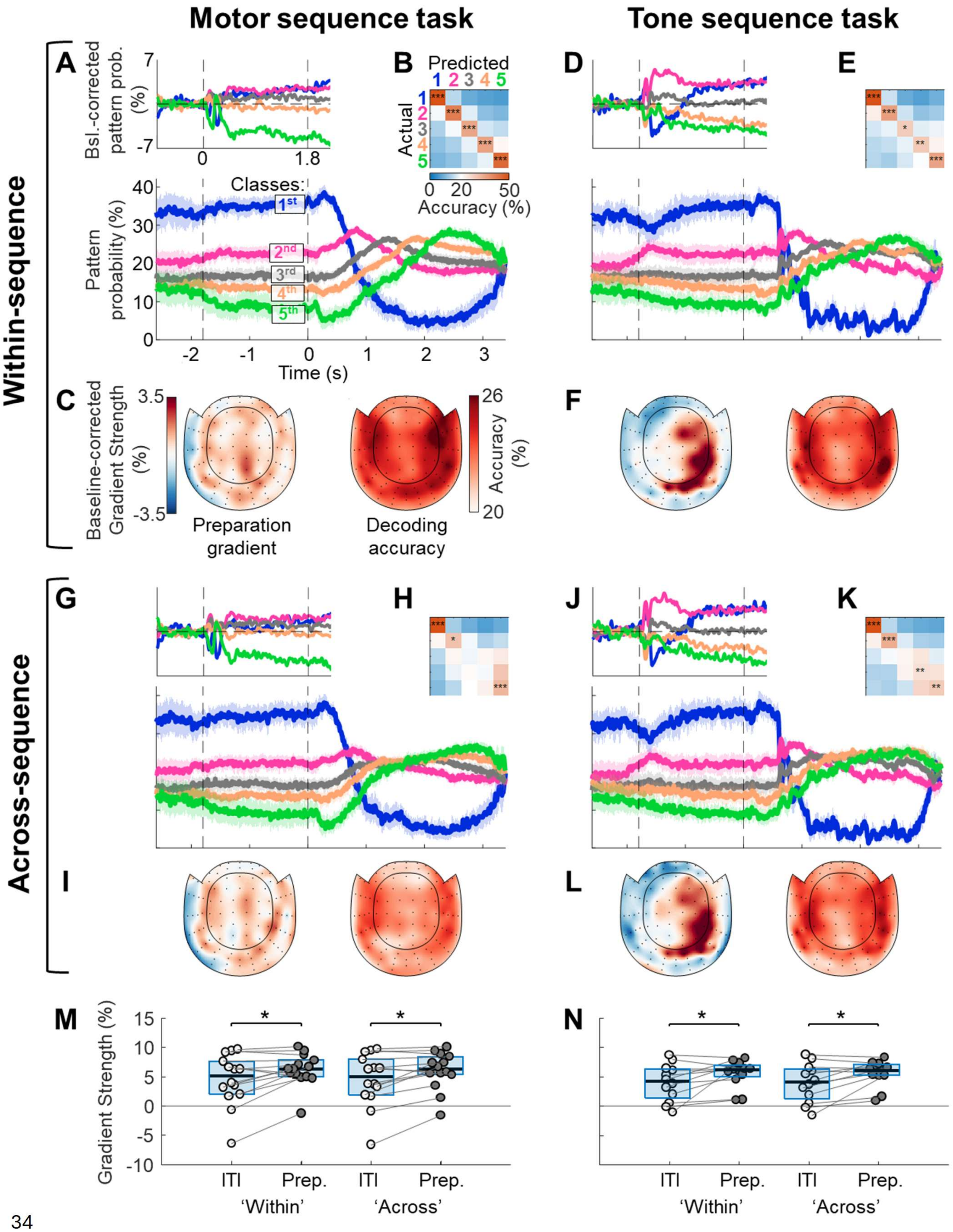


**Fig 2. Results from Experiment 1 MEG data analysis after excluding trials EMG activity.** We ran a control analysis to exclude trials containing unwanted muscle activity. We recorded EMG activity from

the left hand (abductor polices brevis muscle, the abductor digiti minimi muscle, and the first dorsal interosseous muscle with a belly-tendon montage, and the flexor carpi radialis muscle with a belly-belly montage; recorded at sampling rate 1000 Hz, with online band-pass filtering from 0.1 to 330 Hz). To identify which trials contain muscle activity during the preparation time window, raw continuous EMG data were down-sampled to 500 Hz and offline band-pass filtered at 20 to 249 Hz (windowed sinc FIR filter, Hamming window, filter order 826, zero-phase), and then segmented into epochs (-2.6 to 3.4 seconds, time-locked to fractal offset). Then we defined a resting-state noise level from the baseline (ITI) from a copy of the data time-locked to fractal onset using the -800 to 0 ms time window relative to fractal onset. We calculated the signal's RMS per EMG channel from the latest 800 ms before fractal onset ($RMS_{ITI}$). However, since we visually spotted several activity bursts or tonic activity that extends into the ITI, we followed a conservative approach to define the noise level as the lower 10th percentile of the distribution of $RMS_{ITI}$ values over trials, separately per channel ($RMS_{noise}$). In the next step, we calculated the RMS per channel over the entire trial epoch with a sliding 200 ms window to detect also short transient activity. We expressed RMS values at each resulting time point ($RMS_t$) as a ratio relative to the noise level ($RMS_{noise}$) per channel. We set a threshold T = 6, and we flagged trials as to be excluded if they contained a ratio of $RMS_t/RMS_{noise} > T$ for any channel and at any time point of the fractal period (-1.8 which is the latest possible fractal onset to 0 s). Importantly, we did this entire procedure (defining the noise level as the lower 10th percentile and expressing the signal's RMS relative to $RMS_{noise}$ ) separately for each of the two tasks (motor production task or tone sequence task). This is because we observed significant resting-state noise level differences across tasks, which is explained by the different positioning of the left hand (participants operated the response device with their right hand during the tone sequence task, and the left hand, from which EMG activity was recorded, was resting on the side). Another reason is that the motor task's demand to prepare an upcoming sequence through a delay period may have also induced tonic muscle activity as preparation for the imminent execution. Specifically for the tone sequence task, we also flagged trials if they contained an $RMS_t/RMS_{noise}$ ratio higher than T for any channel and at any time point of the listening window (0 to 3.1 s relative to fractal offset). As a last step in our thresholding procedure, any task (motor production or tone sequence task) for which less than 30 trials survived for any of the two sequences, was flagged as not eligible for analysis for this participant. From this analysis, we excluded 5 participants due to a recording artifact that rendered the EMG signal unusable. 14 participants survived for the motor task and 12 participants for the tone sequence task (10 participants overlapping). We performed the same classification analysis as in the main script for the eligible participants/tasks after excluding above-threshold trials. The figure here follows the same structure as **Fig 4** in the main text and shows qualitative similarities as the main analysis. Statistical inference was qualitatively mostly unchanged. Specifically, one-sample tests of Gradient Strength against zero were significant for both decoders, tasks, and trial phases (all $p < 0.001$, Bonferroni-Holm corrected for eight comparisons). A repeated 2x2x2 (within factors Decoder, Task, Phase) ANOVA revealed a significant main effect of Phase ($F_{(1,9)} = 51.260$, $p < .017$, $\eta_p^2 = 0.486$). Planned comparisons for Phase revealed an increase in Gradient Strength from baseline to the last second before cue offset, for both decoders and tasks ('within' decoding, motor sequence task: $t_{(13)} = 2.798$, $p_{corr.} < 0.045$, $d = 0.748$; 'within' tone sequence task: $t_{(11)} = 2.252$, $p_{corr.} < 0.046$, $d = 0.650$; 'across' motor sequence task: $t_{(13)} = 2.927$, $p_{corr.} < 0.048$, $d = 0.782$; 'across' tone sequence task: $t_{(11)} = 2.680$, $p_{corr.} < 0.042$, $d = 0.774$; all $p$-values Bonferroni-Holm corrected for four comparisons). Furthermore, to test whether the EMG thresholding procedure changed Gradient Strength compared to the main non-EMG-thresholded analysis within each participant, we compared the baseline-corrected Gradient Strength between the main analysis and the present supplementary analysis with paired tests per task and decoder. The thresholding did not quantifiably change the baseline-corrected Gradient Strength in either task or decoder (all $t$ <0.99 (absolute values); all $p > 0.975$, Bonferroni-Holm corrected for four comparisons; uncorrected all $p > 0.336$).
